# Reliable, standardized measurements for cell mechanical properties

**DOI:** 10.1101/2023.06.14.544753

**Authors:** Sandra Pérez-Domínguez, Shruti G. Kulkarni, Joanna Pabijan, Kajangi Gnanachandran, Hatice Holuigue, Mar Eroles, Ewelina Lorenc, Massimiliano Berardi, Nelda Antonovaite, Maria Luisa Marini, Javier Lopez Alonso, Lorena Redonto-Morata, Vincent Dupres, Sebastien Janel, Sovon Acharya, Jorge Otero, Daniel Navajas, Kevin Bielawski, Hermann Schillers, Frank Lafont, Felix Rico, Alessandro Podestà, Manfred Radmacher, Małgorzata Lekka

## Abstract

Atomic force microscopy (AFM) has become indispensable for studying biological and medical samples. More than two decades of experiments have revealed that cancer cells are softer than healthy cells (for measured cells cultured on stiff substrates). The softness or, more precisely, the larger deformability of cancer cells, primarily independent of cancer types, could be used as a sensitive marker of pathological changes. The wide application of biomechanics in clinics would require designing instruments with specific calibration, data collection, and analysis procedures. For these reasons, such development is, at present, still very limited, hampering the clinical exploitation of mechanical measurements. Here, we propose a standardized operational protocol (SOP), developed within the EU ITN network Phys2BioMed, which allows the detection of the biomechanical properties of living cancer cells regardless of the nanoindentation instruments used (AFMs and other indenters) and the laboratory involved in the research. We standardized the cell cultures, AFM calibration, measurements, and data analysis. This effort resulted in a step-by-step SOP for cell cultures, instrument calibration, measurements, and data analysis, leading to the concordance of the results (Young’s modulus) measured among the six EU laboratories involved. Our results highlight the importance of the SOP in obtaining a reproducible mechanical characterization of cancer cells and paving the way toward exploiting biomechanics for diagnostic purposes in clinics.

## Introduction

Quantifying the mechanical properties of living cells using an atomic force microscope (AFM^1^) was initiated in the nineties, with the first applications of this technique to soft samples, including living cells^2–5^. Soon, it was reported that cancer cells were softer (i.e., more deformable^5^) than healthy cells (for cells cultured on stiff substrates). Such analogous larger deformability of cancer cells is observed for various cancers such as breast^6^, prostate^7^, ovarian^8^, thyroid^9^, pancreas^10^, and many others^11–14^. In all reported studies, the information on the mechanical properties of cells was derived by applying Hertz-Sneddon contact mechanics^15, 16^ to nanoindentation data. Consequently, Young’s modulus (YM, the proportionality factor between stress and strain) was proposed as a measure of cell mechanics, which is currently in use in virtually all groups worldwide. In parallel with reports on the larger deformability of cancer cells, YM depends on various instrumental and biological factors, making it challenging to obtain the same or even similar values for cells measured in various laboratories using different AFM instruments, possibly under different experimental or environmental conditions. The instrument-related issues stem from deflection sensitivity, spring constant, and tip geometry determinations. Different acquisition settings influence the results, e.g., tip velocity^17^, maximum loading force and the resulting indentation depths^18^, and position on a cell (central part of the cell body or periphery^19, 20^). Details on data analysis, such as how data are processed and which theoretical contact mechanics model is applied, also have an influence. Altogether, these settings affect the final results. Part of the instrumental uncertainties linked to the cantilever spring constant calibration has been elaborated within a previous EU network (COST Action TD1002), leading to the SNAP (Standardized Nanomechanical AFM Protocol) procedure^21^. Using SNAP, a significant reduction of modulus variability during the measurements of living cells can be achieved. However, in this previous study, cells were prepared at one laboratory and sent alive and ready to use by the participating groups. Here, we approached a higher level of complexity in the standardization work toward clinical exploitation since cells had to be cultured locally. Here, we report the results of the effort of the EU ITN *Phys2BioMed* aimed to standardize nanoindentation measurements of mechanical properties of living cancer cells toward clinical exploitation since cells had to be cultured locally. Each participating laboratory applied a standard operational protocol (SOP) while performing the experiments. The SOP includes several steps: cell culturing, AFM sample preparation, nanoindentation measurements, and data analysis. We demonstrated that applying the proposed SOP makes it possible to obtain a similar value of YM for cancer cells, regardless of the nanoindentation instruments (AFM and other indenters; the list of instruments used is included in Supp. Info. List S1) and the location of the measurements. Thereby, we provide evidence that reliable and repetitive assessments of cancer cell mechanical properties in clinical practice are possible. The mechanical properties of cancer cells are of large interest as they can serve as label-free mechanomarkers used, for example, in early-stage diagnosis.

## Results and discussion

Most research has already demonstrated that cancer cells are softer than their healthy counterparts, with Young’s modulus below 10 kPa^6–10, 22, 23^. The larger deformability of cancer cells is typically attributed to a lack of stress fibers in cancer cells and disorganization of the actin cytoskeleton, leading to its larger heterogeneity^6, 9, 24, 25^. The cell line chosen for this study, PANC-1 cells, followed the general trend of actin cytoskeleton disorganization and the absence of thick actin bundles, as commonly observed in cancer cells (Fig. 1).

**Fig. 1.**
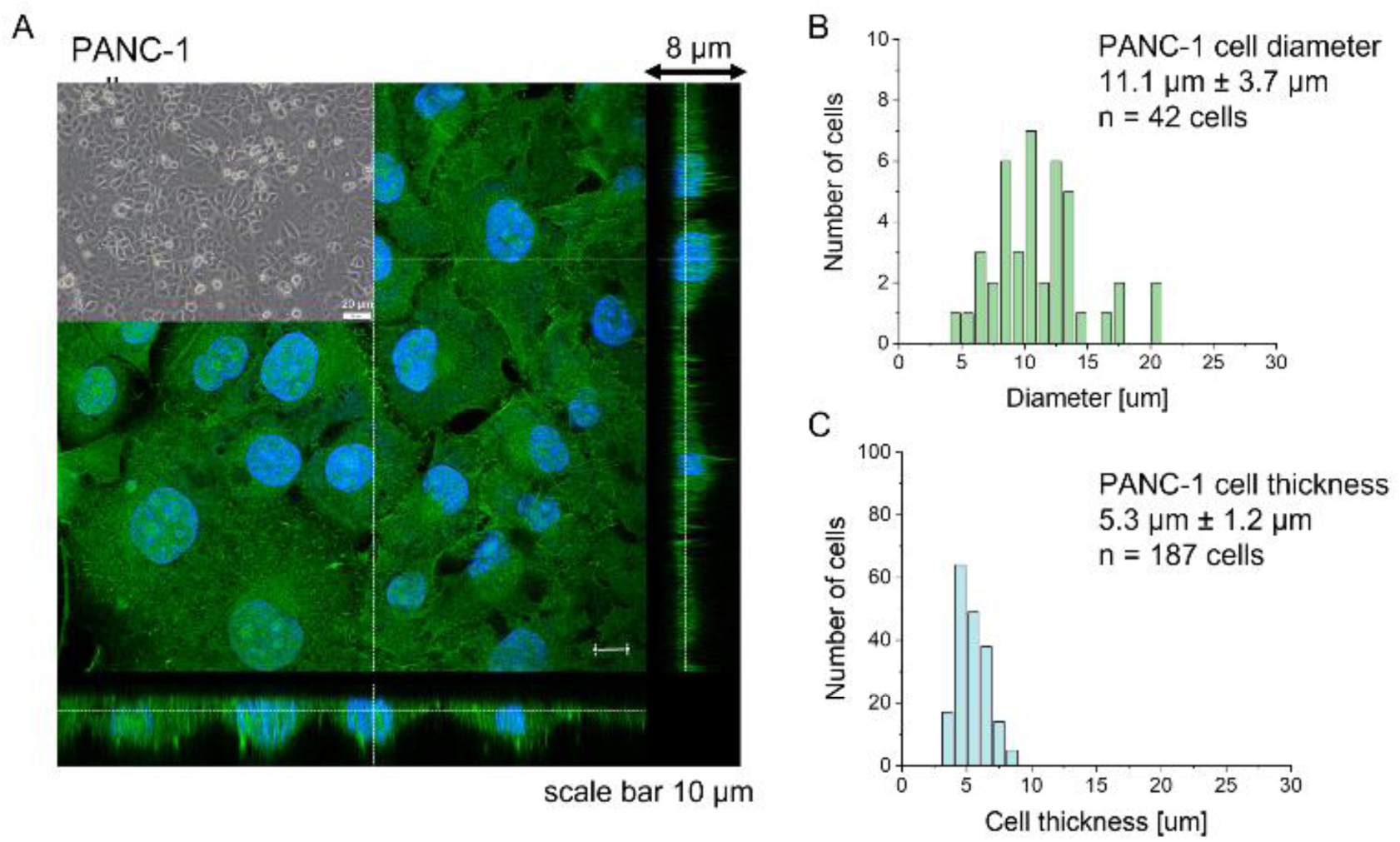
Morphological properties of PANC-1 cells. (A) Confocal images of actin cytoskeleton distribution in PANC-1 cells. F-actin was stained with phalloidin conjugated to Alexa Fluor 488 (green), while cell nuclei were visualized using Hoechst 33342 dye (blue). Inset: a phase-contrast image of PANC-1 cells cultured as a monolayer. From fluorescent images, the cell diameter (B) and thickness (C) were determined as mean ± standard deviation from 42 and 187 cells, respectively.

Analogously to other cancer cells, PANC-1 cells form a monolayer composed of flat cells within which a subpopulation of cells does not adhere to the substrate but forms a secondary layer on top of that attached to the substrate (Fig. 1A, Inset). The nanomechanical characterization was carried out within scan areas of 50 µm × 50 µm encompassing a region of flat cells, meaning that between 3 and 5 cells were probed within a single map. The thickness of the AFM measured cells varied from 3.8 µm to 8.6 µm, depending on whether the measurements were done in the nuclear or pericellular regions. Morphometric analysis of confocal data showed that, on average, PANC-1 cell thickness is 5.3 µm ± 1.2 µm (mean ± standard deviation, s.d., n = 187 cells), while cell diameter is 11.1 µm ± 3.7 µm (n = 47 cells on Fig. 1B&C). In summary, the choice of confluent PANC-1 cells in this study provides cells with a uniformly distributed actin cytoskeleton but also reasonably mimics the heterogeneity of typical cancer cell populations.

The main question of this study is the following: can we obtain similar YM values for PANC-1 cells cultured, prepared, and measured in several different laboratories, following the proposed protocol (Annex 1)? Importantly, the measurements were acquired not only with AFMs but also with other indenters, in which the force signal was acquired differently than in AFM. Such indenters, representing simplified and optimized versions of an AFM, have a likely higher potential to be translated to the clinics. Each laboratory collected at least ten elasticity maps (force volume data) that were analyzed using local software (AFM manufacturers’ software or custom or open-source codes). To design a uniform strategy for handling data analysis and evaluate the software-related variability in the results of PANC-1 mechanics, the elasticity maps were also re-analyzed using one specific custom software at one node of the network (Fig. 2).

**Fig. 2.**
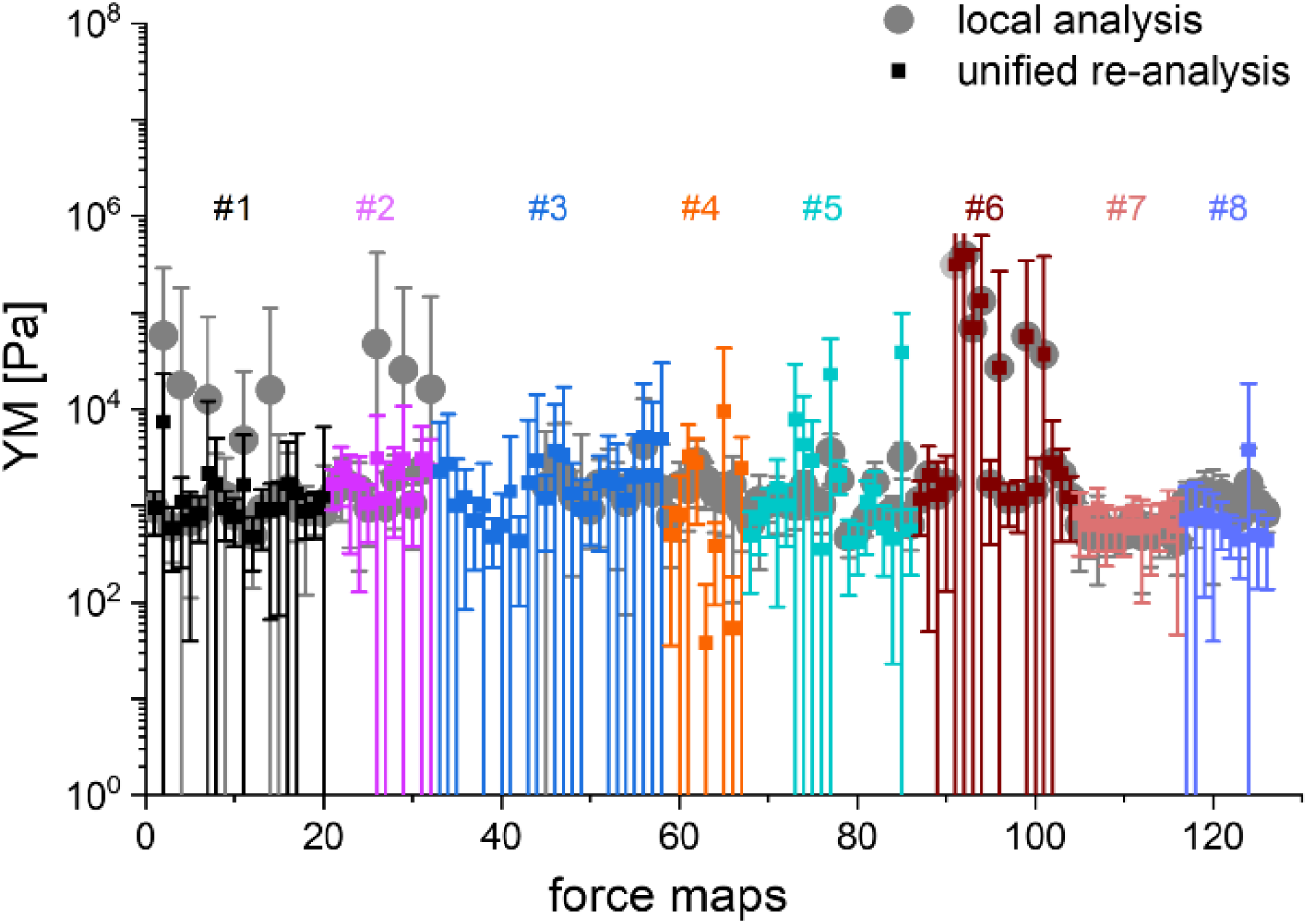
Comparing mean YM values (± standard deviation) obtained by applying local and unified re-analysis to the acquired data of mechanical properties of PANC-1 cells. Each participating laboratory analyzed data using local tools (software provided by the AFM manufacturer or custom codes, solid circles). Next, all data were re-analyzed using the same data processing software (solid squares). The mean and s.d. were calculated and plotted for each recorded map. The color code separates a group of maps recorded using a specific indentation device (mostly AFM) available at the participating laboratory. In total, 125 force maps were recorded in 8 separate experiments.

In most studies, the average YM was used as a measure of cell mechanics (e.g., ref.^26^). Therefore, the mean value of the YM and standard deviation (s.d.) were calculated for each elasticity map. No filtering (e.g., removing bad curves) nor bottom effect correction (BEC,^27, 28^) was applied at this stage. Software-related variability was obtained by comparing the results of the local analysis with the data re-analyzed using the same software. Regardless of the data analysis approach, YM values cluster around 1 kPa, with a mode (maximum peak value) of 1.148 kPa (0.525 .. 2.512 kPa, min, and max YM values) and 1.096 kPa (0.501 .. 2.399 kPa), for the local and the unified analysis, respectively. The statistical analysis (Wilcoxon’s nonparametric test, at a significance level of 0.05) reveals *p* = 0.47562. In most cases, the results overlap; typically, the percentage of change was below 20%. However, in a few cases, the difference between means obtained from local and unified re-analysis was several hundred percent affecting the width of the moduli distribution. The use of unified software for data analysis minimizes this effect. Thus, we conclude that data analysis has little impact on the final YM value if the participating groups follow the same preparation, measurement, and unified data analysis methodology, i.e., the SOP. Moreover, a large standard error (Fig. 2), accompanied by several outlier values, strongly indicates that the YM distributions are not symmetrical. Indeed, the data distribution was characterized by a long tail with high values (Fig. 3A). The YM distribution reasonably follows a lognormal distribution; thus, the modulus is a variable that possesses normal distribution on a logarithmic scale (Fig. 3A, Inset). Despite the clear grouping of YM around 1 kPa, some values can be as large as 300 kPa (Fig. 3A, the x-axis was limited to 10 kPa for better visibility). They frequently appear when force curves are recorded within the peripheral regions of the cell. Their presence affects the calculated mean value leading to an overestimation of the elastic modulus, resulting in a mean of 10.1 kPa and a standard deviation of 46.4 kPa for the acquired data. Such a high mean value and large standard deviation can be explained, besides the intrinsic variability of the cellular system, by the influence of the stiff substrate that can be sensed differently depending on the local thickness of the cell layer, with a stronger effect in the thinner pericellular regions (see discussion below). When data follow lognormal distributions, the median and the 25^th^ and 75^th^ percentiles are better descriptors of cell mechanical properties.

**Fig. 3.**
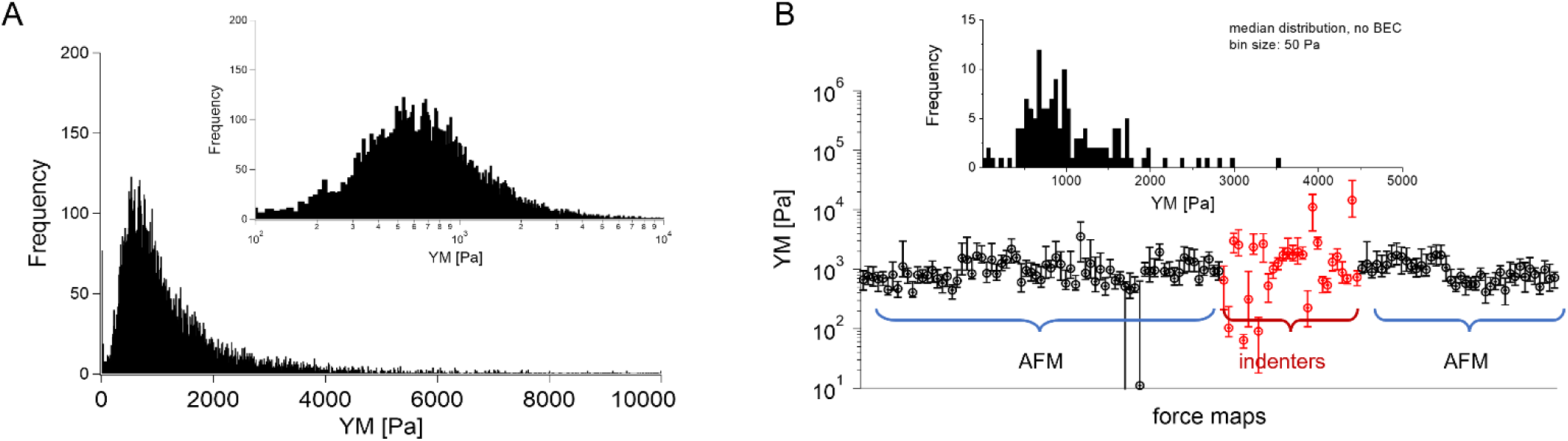
Young’s modulus distributions and median as a descriptor. (A) Lognormal-like distribution of YM values gathered for all recorded force curves (14 000 force curves) using unified re-analysis. Inset: The histogram is approximately symmetric when represented on a logarithmic scale, a signature of log normality. (B) Median YM values (error bars: +75th and –25th percentiles) plotted for each recorded map (medians were calculated for the data presented in Fig. 2, the measurement technique is indicated and marked by colors). Inset: Distribution of the median YM values, from all maps recorded.

The median is less affected by very large YM values, e.g., outliers because of bad force curves (e.g., because the tip does not fully retract due to adhesion), than the mean. The median values determined for each force map recorded for cancer PANC-1 cells vary around 1 kPa, too (Fig. 3B, where error bars enclose the 25^th^-75^th^ percentile range). Simultaneously, medians reveal smaller elastic modulus variability within a single map. However, the overall mean median value, calculated for all recorded maps, equaled 1.196 kPa ± 1.524 kPa (s.d.). It should also be noted that the medians determined for the individual maps are normally distributed in a logarithmic scale (Fig. 3B, Inset). A large standard deviation indicates the intrinsic variability present in the specific experiment. The presence of markedly different YM values in the recorded data sets is not surprising because the size of each elasticity map was set to 50 µm × 50 µm (the corresponding grid of 10 pixels × 10 pixels was set). The diameter of a single PANC-1 cell was around ∼ 11 µm (see Fig. 1); thus, a few cells can be observed within a single map (as shown in the exemplary map presented in Fig. 4A,B). The higher parts show regions around the cell nucleus, while the thinner regions are the cell periphery. Three cells were probed within the map (marked as #1, #2, and #3 in Fig. 4A).

**Fig. 4.**
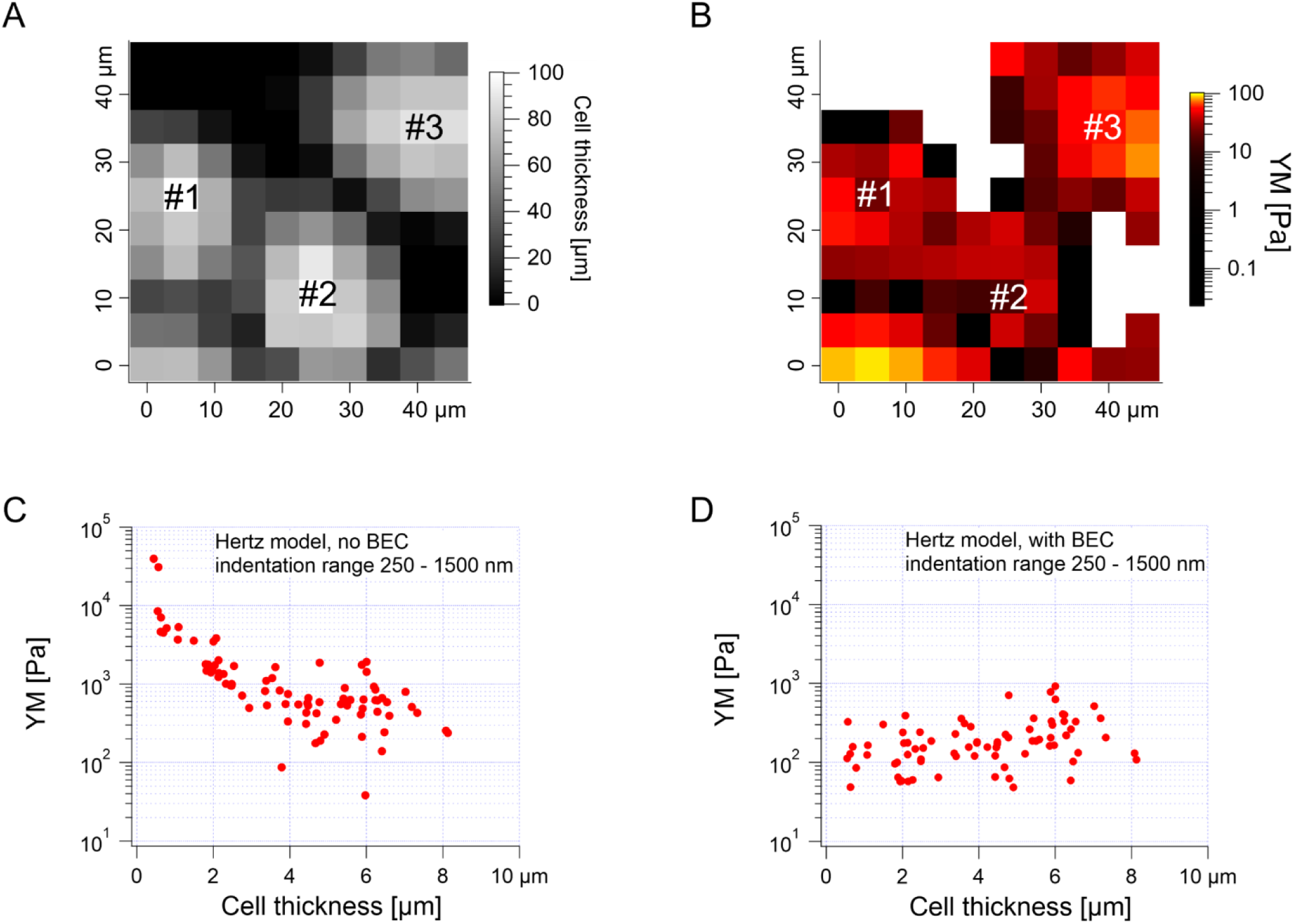
High YM corresponds to the lowest cell thickness, indicating substrate influence. (A) An exemplary map showing the variation in the local cell thickness. Three cells were measured within the probed area. (B) Exemplary elasticity map recorded for the PANC-1 cell monolayer revealing regions with low (up to 10 kPa) and high (above 10 kPa) modulus. A high YM, marked by white squares, contributed to the presence of outliers. (C, D) YM plotted as a function of cell thickness, before (C) and after (D) bottom effect correction (standard BEC approach).

The darker topographic areas correspond either to very thin regions of the cells or to the exposed substrate. A steep slope characterized the force curves recorded within the darkest regions, similar to the slope measured in the contact region of the calibration curves, confirming that the stiff underlying substrate is present within this map (data not shown). Usually, the measured Young’s modulus was higher when moving towards the cell periphery than in the middle of the cells (above or near the cell nucleus), i.e., along a path of decreasing cell thickness. The higher YM values, measured in the thinner regions, particularly those measured at the cell–exposed substrate border, will significantly impact the YM mean, shifting it towards higher values. Fig. 4C shows a strong correlation between the measured YM and local cell thickness, suggesting that a bottom effect correction (BEC) must be applied to the force curves^27, 29–31^. This strong bottom effect can be understood by considering that we used large colloidal probes in our experiments. Indeed, BEC is implemented by multiplying the point by point of the force curve by a polynomial function *Δ* of the non-dimensional parameter *χ*, which represents the contact radius ratio to the cell thickness (Eqs 2-4).

When using sharp tips on relatively thick samples, the corrective function *Δ* is close to unity, *χ* is close to zero, and; in our case, however, hemispherical tips with a radius of 5.5 µm were used during the indentation experiments, which boosted the impact of the bottom effect, especially in the thinner regions of the cell layer. Therefore, the application of BEC to the data is necessary. Fig. 4D confirms the efficacy of the bottom effect correction, which removes the YM dependence on the cell thickness. When working on confluent cell layers, the substrate may not be accessible in topographic maps, which limits the application of the BEC; indeed, the standard BEC approach requires the exact knowledge of the local cell thickness, which can only be measured relative to the substrate. In our case, the substrate was recorded only in 22 out of 141 maps acquired in different experiments conducted with different AFM instruments at different locations. The situation can be even worse, for example, when the cell layer’s confluence is complete. Since the bottom effect can be important, especially when using large colloidal probes, we developed an approach to be applied in conditions when a reference substrate for the accurate determination of the local thickness of the sample is totally or partially missing. We will refer to our approach as the estimated BEC method. The estimated BEC approach we proposed in this work relies on the determination of the local cell thickness by means of confocal microscopy to determine the mean thickness of the cell layer, and AFM topographic maps, to determine the relative thickness variations around the mean value. By adding the relative thickness variations to the mean thickness of the cell layer, we obtained the estimated local cell thickness (see Methods for details). When a few topographic maps with an exposed substrate exist, these maps can be used to estimate the mean thickness of the whole cell layer, which is supposed to be reasonably accurate as long as the number of imaged cells is sufficiently large. In our case, the average (for different cells) of the maximum cell thickness was estimated to be 5.3 µm using confocal microscopy (Fig. 1, 187 cells). From the 22 topographic AFM maps with the exposed substrate, the cell height was estimated to be 9.1 ± 1.9 µm. These much larger values stem from various reasons, mainly from the distinct cell preparations (fixation and staining for confocal microscopy versus living cells for AFM) and contact point determination used to calculate the cell height. In the case of spherical probes, surface glycocalyx or membrane corrugations may influence the determined cell height because the cantilever deflection reflects the interaction between the probe and the cell surface. The reported size of the surface brush for cancer cells can reach even a few microns in length^32^.

We first tested the accuracy of the estimated BEC approach on the 22 maps with the exposed substrate since, in this case, the local cell thickness can be accurately measured, and the standard BEC can be applied accurately. Notably, these 22 maps were recorded in 4 different experiments (location, AFM instruments); therefore, we applied standard and estimated BEC to data from each experiment separately (labeled as Exp 1 to 4, Fig. 5). The YM distribution without the BEC (Fig. 5A) showed data centered around 1 kPa with a long right tail reaching 100 kPa. Distributions were consistent among the four experiments. After applying the standard and the estimated BEC, all moduli were shifted towards lower values, with a median in the range of 100 – 200 Pa. The standard and the estimated BEC methods were similar regarding the overall shape, median values, and median absolute deviations of the YM distributions.

**Fig. 5.**
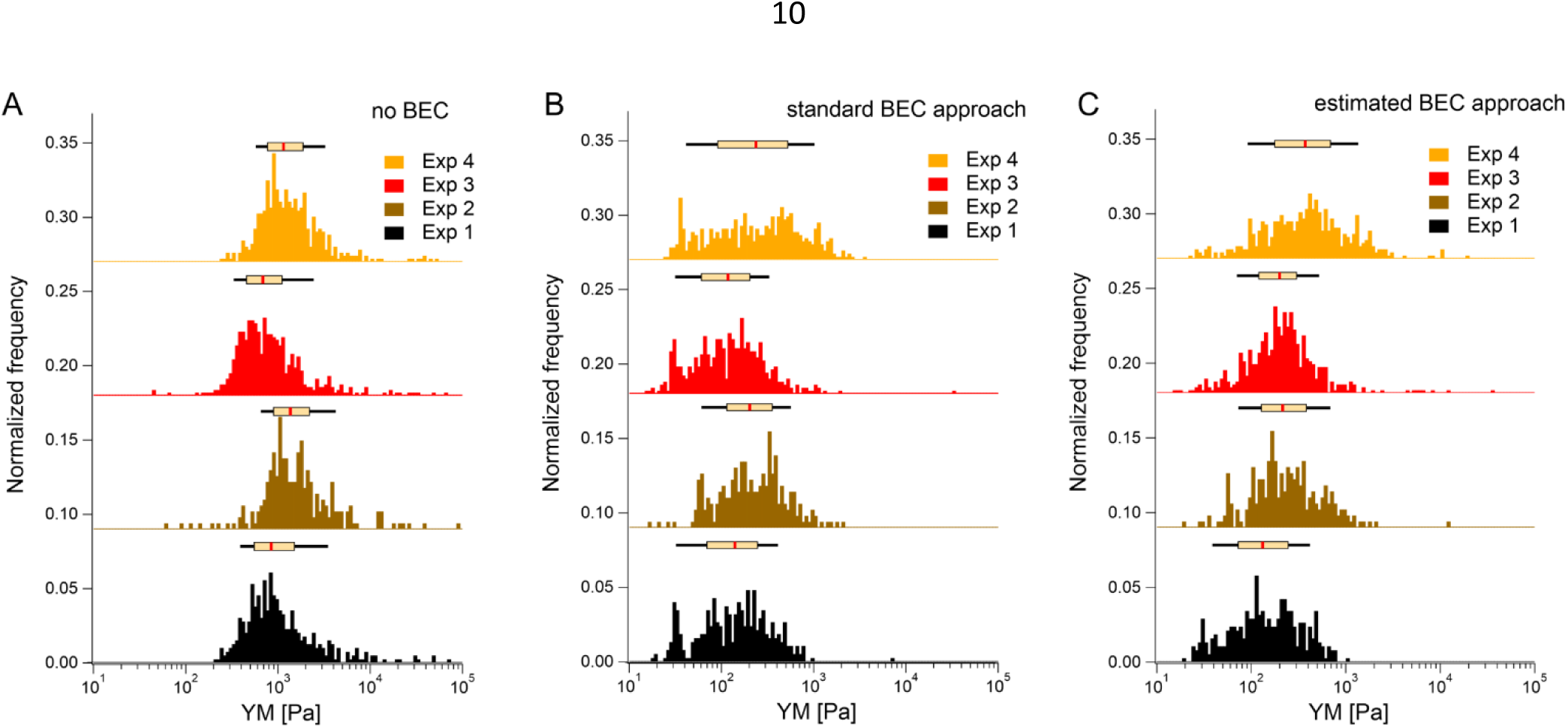
Bottom effect correction (BEC) demonstrated on datasets originating from 4 distinct experiments. Each experiment reflected several force maps recorded from one sample in one laboratory, pooled into one histogram. In total, 22 maps were recorded, where the cell thickness could be accurately determined from the AFM data. (A) Modulus distribution without BEC. (B,C) The corresponding histogram after BEC considers the exact (B, standard BEC) and approximated (C, estimated BEC) cell thickness (here, 5.3 µm, obtained from confocal microscopy). Histograms of all YM (each force curve is considered) were pooled together depending on the analysis approach, i.e., no BEC, standard, and estimated BEC approaches. The box plot on top of each histogram refers to the median, 25^th^ & 75^th^ percentiles (box), and 10^th^ & 90^th^ percentiles (black line). Note: the baselines were shifted in the Y direction to separate histograms.

Having demonstrated the validity of the estimated BEC method based on the estimation of the local cell thickness in the absence of the reference substrate, we applied it to all available maps collected network-wise, in most of which the substrate was not visible in the topographic maps (Fig. 6).

**Fig. 6.**
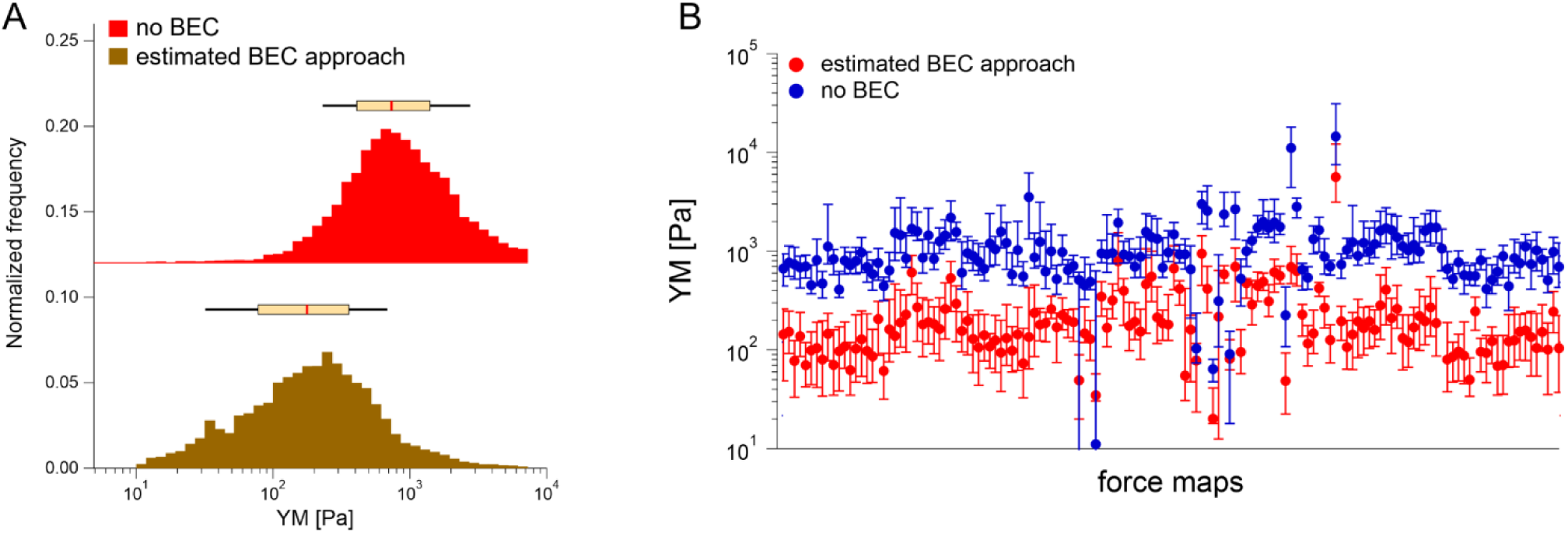
The results of the estimated BEC applied to all data where the cell thickness cannot be directly measured by AFM. (A) The maximum modulus distribution was found between 100 – 200 Pa. YM histograms (each force curve is considered) are pooled together depending on the analysis approach, i.e., no BEC or the estimated BEC approach. The box plot on top of each histogram refers to the median, the 25^th^ and 75^th^ percentiles (box), and the 10^th^ and 90^th^ percentiles (black line). (B) The corresponding medians without and with the estimated BEC approach plotted for each recorded map (error bars: +75^th^ and -25^th^ percentiles). Note: the baseline was shifted in the Y direction to separate both histograms Finally, it is interesting how the proposed approach of the standardization experiments relates to a more common approach (Fig. 7).

**Fig. 7.**
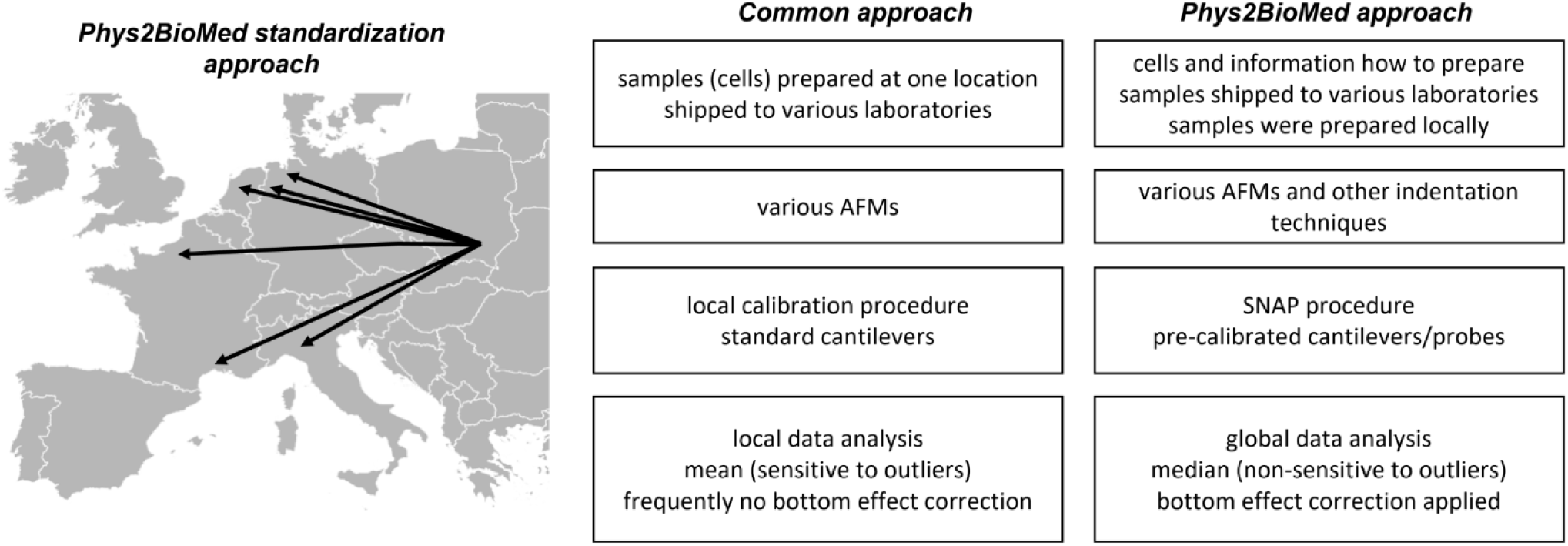
Phys2BioMed Standardized Operational Procedure (SOP) for measuring mechanical properties of cells. The advantages of the proposed Phys2BioMed standardization approach were compared to the more common approach.

YM distributions before and after the estimated BEC correction are shown. The distributions peaked at approximately 0.700 kPa and 0.170 kPa, respectively. The medians calculated for each recorded map before and after using the estimated BEC approach are presented in Fig. 6B. The average medians are 1.196 kPa ± 1.524 kPa (mean median value ± standard deviation of the medians) and 0.253 kPa ± 0.489 kPa before and after correction, respectively. Interestingly, the variability of medians remained at the same level, regardless of whether the estimated BEC was applied or not. As medians are less sensitive to outliers and BEC correction eliminates the influence of the stiff underlying substrate, the observed variability indicates intrinsic heterogeneity of PANC-1 cancer cells.

The more common approach for the standardization of measurements uses the samples (also living cells^21^) prepared in one laboratory, followed by their shipment to other laboratories participating in the standardization process. Then, the samples were measured using various instruments with cantilevers calibrated using local software using the thermal method. The data are analyzed, most frequently, using locally available tools; however, as shown during the development of the SNAP procedure, the analysis of data recorded by various groups applying SNAP eliminates variability linked with deflection sensitivity calibration^21^. Typically, the final results are presented as the mean (average) value, whereas variability is assessed by standard deviation.

## Experimental

### Cell lines

A human pancreatic cancer cell line, PANC-1 (ATCC, CRL-1469, LGC Standards, RRID: CVCL 0480), was chosen for the measurements. These cells were isolated from a pancreatic carcinoma of ductal cell origin derived from the tissue of a 56-year-old male. Cells possess an epithelial-like morphology, and they are adherent. PANC-1 cells were cultured in DMEM supplemented with 10% FBS and 1% antibiotics (a detailed procedure for handling the cells for standardized AFM measurements is presented in Annex 1).

### Shipping the cells

PANC-1 cells were cultured in the plastic culture flasks with a surface area of 25 cm^2^ (Saarsted) up to 60-70% of their confluency in the culture medium (the term culture medium refers here to DMEM supplemented with 10% of FBS and 1% of antibiotics). Cell culture was conducted in the CO_2_ incubator providing a temperature of 37°C and an atmosphere of 5% CO_2_ and 95% air. Then the culture flask with cells was removed from the incubator. Next, the medium in the culture flasks was replaced with the fresh one, fully filling the flask (up to the stopper). Flasks were closed and moved from sterile condition to a foam box (RT) prepared to be sent to participating laboratories. The parcels were shipped to other countries with the next-day delivery order till noon; however, the shipping lasted up to 2 days. Nevertheless, each participating group received cells in good shape. Before shipping, cells were mycoplasma tested (see Annex 1).

### Sample preparation

To standardize the AFM measurements, a timeline of the experiments was prepared (a detailed procedure is described in Annex 1). Briefly, after the arrival, cells were passaged onto a Petri dish in the following way. Cells were washed with 1 mL of PBS (Phosphate Buffered Saline, Sigma) solution. Next, 2 mL of pre-warmed 0.25% trypsin/PBS solution was added to the culture flask for 3 min in the CO_2_ incubator. When more than 90% of cells were detached, they were transferred to a 15 mL tube, to which 3 mL of culture medium was added. Cells were centrifuged at 1800 rpm for 4 minutes. After removing the supernatant, a 2 mL fresh culture medium was added, and cells were gently aspirated with a pipette to obtain homogenous cell suspension. The number of cells was counted using the Brürker chamber, and the suspension was diluted at the final concentration of 150 000 cells/mL. Then, 2 mL of cell suspension was moved to a Petri dish (TPP, with a surface area of 9.4 cm^2^, the number of cells was adjusted according to the different sizes of the Petri dish when needed; therefore, the cell seeding density was the suitable parameter.). After adding 4 mL of culture medium, the Petri dish with cells was placed into the CO2 incubator for 48h prior to AFM measurements. AFM measurements were conducted on a cell monolayer, at 37°C, in DMEM supplemented with 10% FBS, 1% antibiotics, and 10 mM of HEPES (Sigma).

### Cantilevers

MLCT-SPH-DC silicon nitride cantilevers (hemispherical version of Bruker MLCT-BIO-DC probes) with a hemispherical tip of a 5.5 µm radius were chosen (Supp. Fig. S1). The manufacturer pre-calibrated the spring constant of these cantilevers using a Laser Doppler Vibrometer; its value varies between (140 mN/m and 220 mN/m). In total, eight pre-calibrated cantilevers were used in this study. The choice of the cantilevers was dictated by the well-defined spherical shape of the probing tip and by its size, which minimizes non-linearities and provides good averaging of the mechanical data; moreover, pre-calibration of the spring constant eliminates instrument-dependent variability in the calibration process. Moreover, contactless SNAP protocol 21 was applied to calibrate the deflection signal sensitivity. For the nanoindentation measurements, cantilevers (Optics11 B.V.) with 2.5-3.2 µm radius and 0.021-0.025 N/m were used. The spring constant was calibrated following the procedure detailed in Beekmans et al.^33^, using a weighting scale with 100 ng sensitivity (MSA2.7S-000-DF, Sartorius AG).

### AFM and indentation measurements

AFM-based elasticity measurements were performed in six laboratories with different instruments (see Supp. Info. List S1).

Measurements were conducted on the flat part of the PANC-1 cell monolayers (Supp. Fig. S2). The AFM tip was moved over a flat part of the monolayer. Ten elasticity maps of 50 µm × 50 µm were acquired. Within the recorded area, a few cells can be visualized. A grid of 10 pixels × 10 pixels was set on each map. The other parameters set during the experiments were: z-travel distance 5 µm, number of points in the force curve: possibly 1 point/nm (i.e., n > 5000), cantilever approach velocity of 5 µm/s, force scan rate of 0.5 curves per second, trigger point (the point where approach ends) of 700 nm or 7 nN, z-close loop – ON. For nanoindentation measurements, the same area scan parameters were set. The experiments were performed in displacement control mode, with a z-travel of 5 µm over 1 s. The sampling rate was set to 1 kHz.

### Data analysis

The mechanical properties of PANC-1 cells were quantified using the Hertz-Sneddon contact mechanics^15, 16^, delivering Young’s modulus. Briefly, the force curves were recorded on a stiff, non-deformable surface (calibration curve recorded here on a Petri dish surface) and on cells. The approach part of the force curve is considered here. Next, the calibration curves were subtracted from those recorded on cells. The obtained force versus indentation curves were fitted with the equation relating the load force F and indentation δ for the spherical probe:

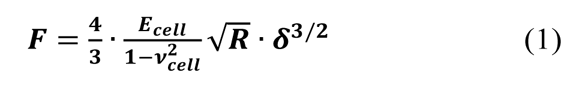

where *R* is the radius of the AFM probe, *ν_cell_* is the Poisson ratio of the cell (set to 0.5, treating cells as incompressible materials), and *E_cell_* is Young’s modulus, which was determined for the whole indentation range. The indentation range considered for the Hertzian fit was 250-1500 nm. Importantly, used hemispherical probes with a radius of 5.5 µm fulfill the Hertz model requirement of a small enough contact area between the probe and cell surface. The data were analyzed locally using various accessible software and re-analyzed using Igor Pro-based procedures (Wavemetrics, Lake Oswego, OR, USA).

### Bottom effect correction

The effect of a stiff underlying surface on the mechanical properties of cells is more pronounced for spherical probes indenting the cell surface^27, 29, 31^. Thus, the following equation was used to correct the force curve (as described in the section):

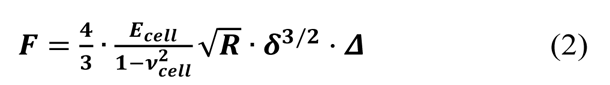

where *F* is the force, *E_cell_* is the elastic (Young’s) modulus of the cell, *R* is the radius of the indenting spherical probe, and *δ* is the indentation.

The correction factor *Δ*, for bonded samples^27^ (since we measured cell monolayer), is defined as follows:

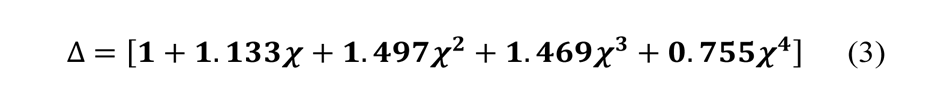

where *χ* depends on *R* and *δ* as:

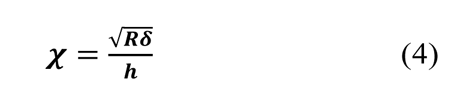

where *√Rδ* is the contact radius, and *h* is the (local) cell thickness. The correction factor *Δ* is, therefore, a function of indentation *δ*, which must be applied point by point to the experimental force curve, and depends parametrically on both the tip radius *R* and the local cell thickness *h*.

### Determination of the topographic

The cell local thickness was determined by analyzing the contact point within each force curve, a fitting parameter of the Hertzian fit. Force curves taken on the substrate were selected based on the criterion that the slope in the contact region is larger than 0.9. Theoretically, calibrated force curves (cantilever deflection vs. z-piezo displacement) measured on a stiff substrate should have a slope in the contact region of 1. However, due to noise in the instrument or adsorbed molecules from the medium on the support, we often see a smaller slope, depending on the loading force. Thus, using a criterion based on the slope at medium forces, we proved to successfully select only force curves on the substrate. These contact points define a plane, which determines the substrate. This substrate plane is subtracted from all contact points (including those taken on the cell), resulting in the local cell thickness. Then a second fit of the force vs. indentation data using the BEC model considering the local cell height is performed to get “true” Young’s modulus value.

### The estimated BEC method

The standard bottom effect correction procedure described above requires that the local cell height (or cell thickness) be known for each force curve. However, this can be done only if there are regions of the substrate in the corresponding topographic map (obtained as described in the previous section); this is not the case, for example, for confluent cell layers. Thus we have designed the following approximate procedure to estimate the local cell height in case the substrate is not accessible in the maps.

First, the typical cell height (i.e., the maximum thickness of a particular cell) for a certain number of cells is determined by other techniques, e.g., by confocal microscopy (if possible and enough maps are available, this can also be calculated from the subset of AFM topographic maps where the exposed substrate is present). The average cell height (exactly the average height, i.e., the maximum thickness, over several cells) is then used as the typical cell height. When analyzing force maps with no substrate data, we assign a thickness according to the typical cell height to the maximum contact point in this force map. Since we occasionally have extreme outliers due to bad force curves, we do not take the maximum contact point but take the 95^th^ percentile in the distribution of the contact points as the “representative” maximum value. The other force curves will be assigned a thickness accordingly. This thickness value is then used in the BEC. Sometimes this algorithm leads to negative thickness values since the range of topography in this particular force map is larger than the typical cell height. In this case, thickness values are corrected by adding a constant so that the smallest thickness becomes zero.

### Statistics

As stated in the text, results are expressed as a mean ± standard deviation or median ± standard deviation of the medians. Statistical significance was determined using nonparametric Wilcoxon’s test at a statistical significance of 0.05.

## Conclusions

Here, we elaborated on the standard protocol for culturing cells at each participating laboratory (Annex 1). The mechanical properties of single cells, even if they originate from the same cell line, are not uniform due to the variability in cytoskeleton organization, shape, substrate properties, etc.^34–36^. This, together with differences in cell cultures, preparing them for indentation measurements and measurement conditions, contribute to the relativeness of YM and its large variability observed in various already reported data^34–36^. Single cells lack intercellular connections moving them far from physiological conditions. Most cancer cells tend to grow as clusters, already at the beginning of cell culture. Although the stock of cells was shipped from one laboratory to other participating laboratories as living cells in culture flasks, we found that the delivery time was less important than the time of the measurements after passaging cells. Once cells were passaged (directly after arrival), AFM measurements were conducted after 48 h in this study. Various AFM instruments and other indenter techniques were applied to measure the mechanical properties of living PANC-1 cells using pre-calibrated cantilevers (known spring constants) with hemispherical probes. The data were analyzed using the same software, eliminating the problem of various algorithms, e.g., the most critical here is the determination algorithm to determine the contact point position. The analysis revealed that the median is a better descriptor of the mechanical properties than the mean, as it is less sensitive to the presence of outliers and it better handles lognormal distributions. Applying BEC correlates Young’s modulus to the cell thickness and therefore minimizes the influence of the stiff substrate. We demonstrate that the determined cell thickness based on confocal images can be used in BEC calculations, despite its obvious inaccuracy. In conclusion, we demonstrated that standardizing indentation measurements of cell mechanical properties using SOP enables obtaining similar results among various laboratories, opening the door for robust clinical applications.

## Author Contributions

SPD, SGK, KG, HH, ME, EL, MB, NA, MLM, JLA, LRM, VD, SJ, SA, KB, FL, HS, FR, AP, MR, ML contributed to the development of the protocol, designing the study, local measurements and data analysis, interpretation, and manuscript preparation; SPD, KG, MR re-analyzed data using unified software; HH contributed to development of the estimated BEC method; JP was responsible for cells preparation and shipping; JO, DN contributed to protocol development; ML, AP, MR wrote a draft. All authors contributed to manuscript editing and revising.

## Supporting information

Supporting Materials with Annex 1

## Acknowledgements

We acknowledge the support of the European Union’s Horizon 2020 research and innovation programme under the Marie Skłodowska-Curie grant agreement No. 812772, project Phys2BioMed. We would like to thank Alexander Dulebo (Bruker) for his advice on hemispherical AFM probes. We would like to thank Massimo Alfano and Laura Vidal for contributing to the discussions during the Phys2BioMed meetings and Matteo Chighizola for support of the BEC method.

## Supporting Information

**Supporting Figure S1.**
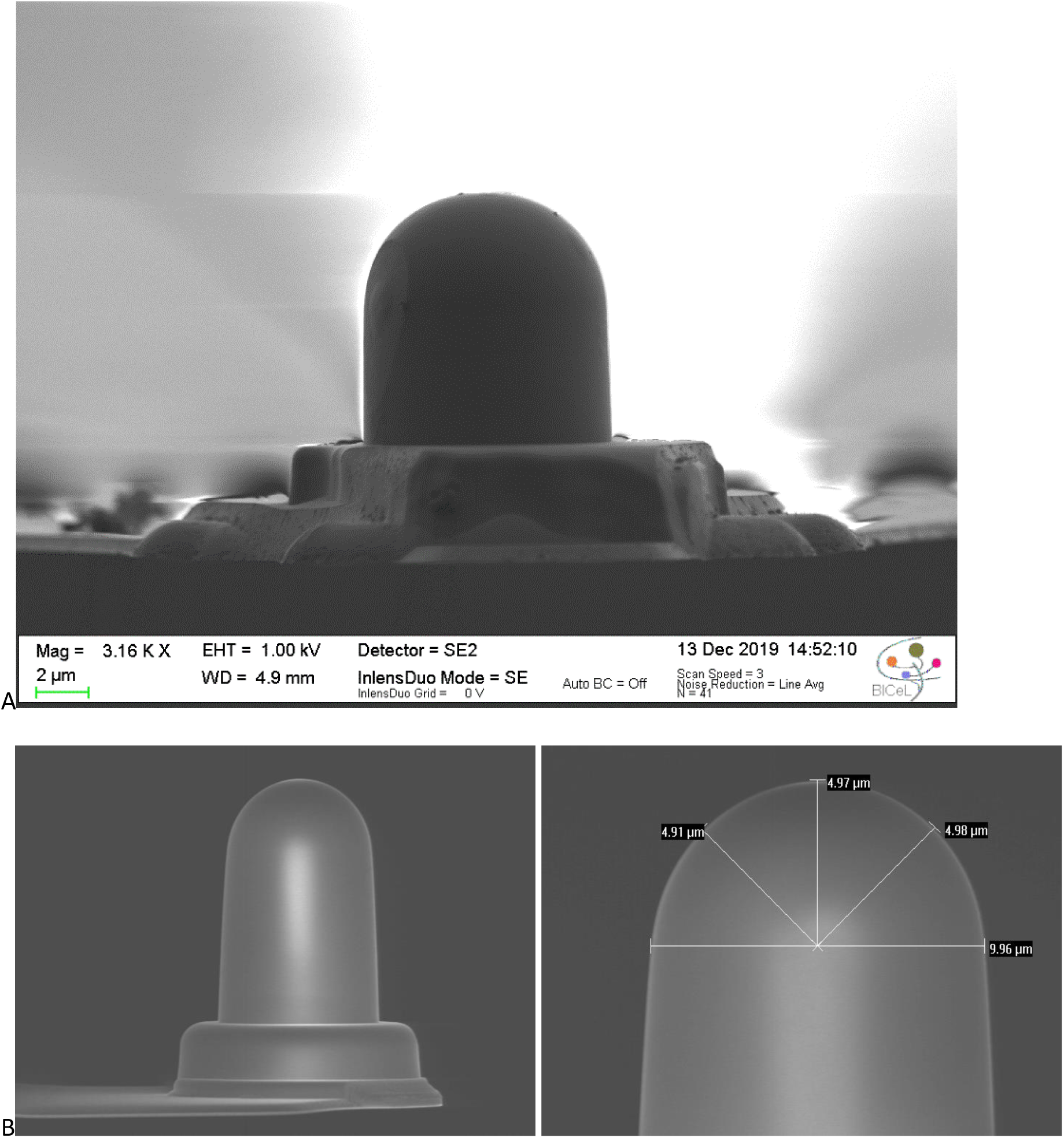
**A)** SEM (Scanning Electron Microscope) image of the MLCT-SPH-DC silicon nitride cantilevers (Bruker) with a hemispherical tip. The image was collected using SEM available in BICeL facility (Lille, France). B) SEM images of the currently available cantilevers with SPH tips available for AFM community (courtesy of A.Dulebo, Bruker).

**Supporting Figure S2.**
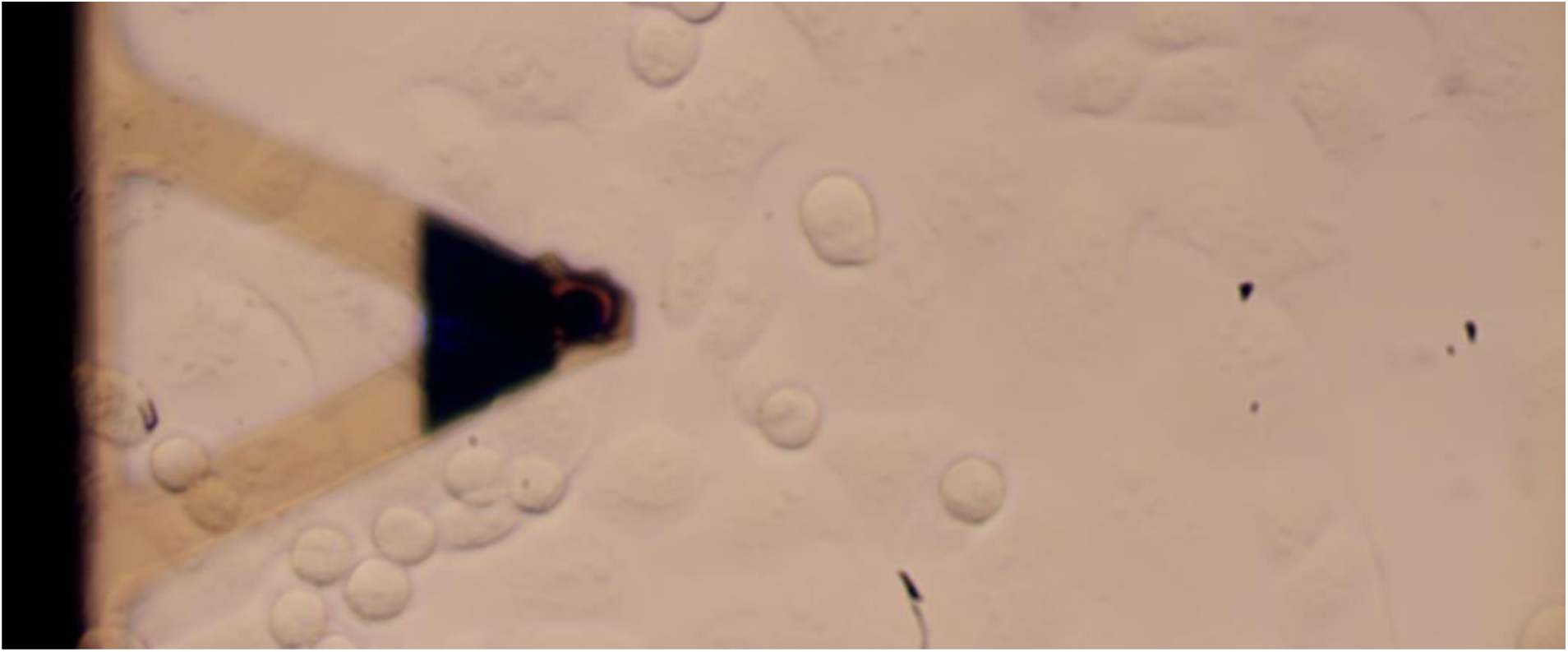
**Supporting Figure S2.** Top view image of the PANC-1 monolayer, the flat area of the monolayer measured in standardization experiments. Image was collected with the top view optics integrated with MFP3D (Asylum Research, Santa Barbara, CA, USA).

## Annex 1: Protocol for measuring cell mechanics

### Protocol sheet for measuring cell mechanics

#### 0. General Information

Materials needed:

- two 25 cm^2^ culture flasks (#1, #2) with living cells (here, the protocol was tested for pancreatic cancer PANC-1 cells);

**Note:** PANC-1 cells were established from a human pancreatic cancer, isolated from a pancreatic carcinoma of ductal cell origin of a 56-year-old male (Lieber et al. International Journal of Cancer. 15 (1975) 741–747, doi:10.1002/ijc.2910150505). These cells have an epithelial morphology and are adherent in cell culture flasks; however, they tend to clump. PANC-1 cells can metastasize, but they have poor differentiation abilities.

- four empty culture flasks (sterile)
- two 50 ml tubes of culture medium (DMEM + 10% FBS) with antibiotics (1%) - (sterile);
- one 50 ml tube of PBS (sterile)
- one 15 ml tube with trypsin solution (sterile);
- one 15 ml tube of freezing medium: DMEM + 5 % DMSO (sterile)
- one 2 ml tube of 1 M HEPES (sterile)

Cell Culture Dish, 35×10mm – TPP cat no. *93040*

Tissue Culture Flasks 25 cm2 – TPP cat no. 90025

DMEM - ATCC cat no. 30-2002 FBS - SIGMA cat no. F9665

Trypsin-EDTA solution (10x) - SIGMA cat no. T4174 Dimethyl sulfoxide (DMSO) – SIGMA cat no. D2438 HEPES buffer (1 M) – SIGMA cat no. H0887

PBS – SIGMA cat no. P4417

Antibiotics (Penicillin − Streptomycin − Neomycin Solution Stabilized) – SIGMA, cat no P4083

#### 1. Shipment of cells

Living cells can be shipped at 50-60% confluency; culture flasks should be filled with a maximum amount of culture medium (for PANC-1 cells: DMEM + 10% FBS + 1% antibiotics).

1. Document sample and shipping details:

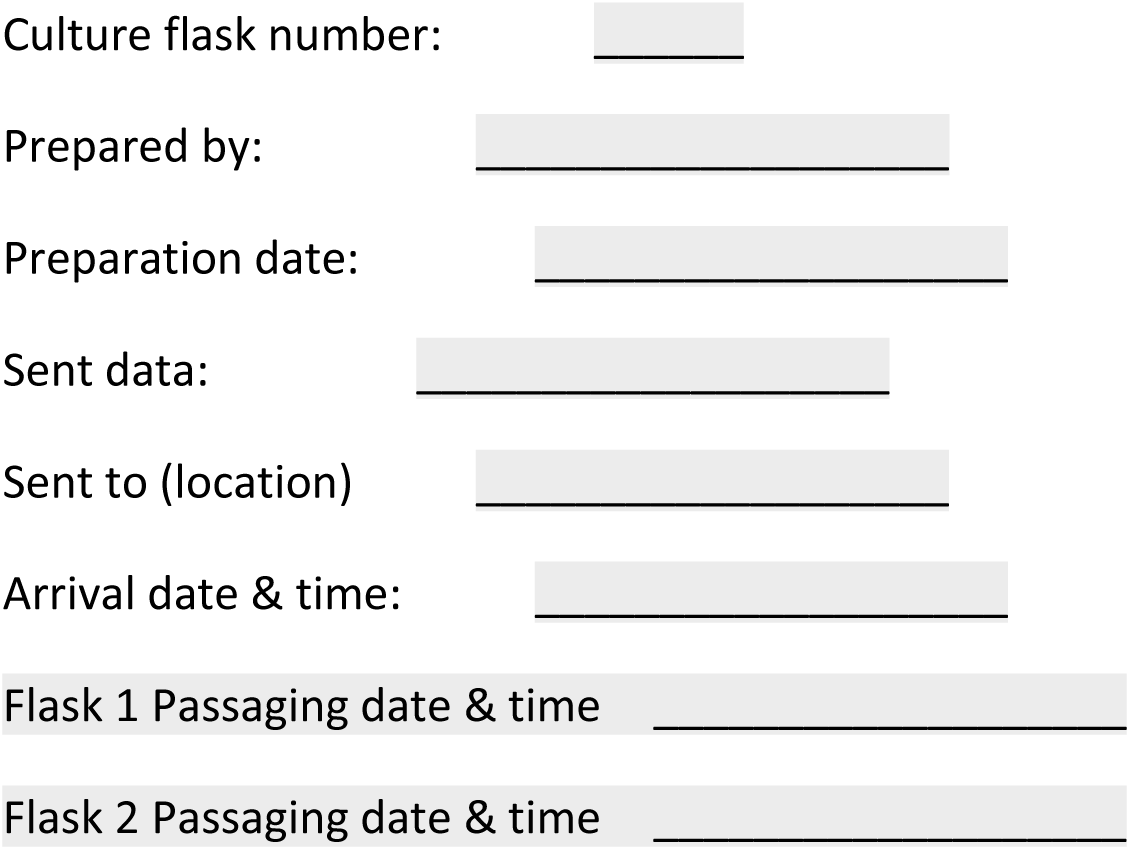

#### **2.** Cell culture

Work under sterile conditions (under the laminar flow chamber):

1. After arrival, remove half of the culture medium and exchange the bottle stopper (take a new one from an empty culture flask)
2. Take the culture flasks and make a passage according to cell **procedure #1**.
3. Leave cells **overnight** in a CO2 incubator.

**Protocol #1 (cell passaging, preparation of cells for AFM measurements):**

1. Remove and discard the culture medium.
2. Briefly rinse the cell layer with 1 mL PBS solution to remove all serum traces containing trypsin inhibitor.
3. Add 2.0 ml of pre-warmed 0.25% Trypsin/PBS solution to the culture flask and place the flask for 3 minutes in the CO2 incubator. Observe cells under an inverted microscope until the cell layer is dispersed (usually, it takes 5 to 15 minutes).

**Note:** To avoid clumping, do not agitate the cells by hitting or shaking the flask while waiting for the cells to detach.

4. When ≥ 90% of the cells have detached, tilt the vessel for a minimal time to allow the cells to settle.
5. Transfer the cells to a 15-ml conical tube, add 3mL of culture medium (DMEM +10% FBS + 1% antibiotics), and centrifuge them at 1800 rpm for 4 minutes.
6. Remove the supernatant – a cell pellet is located at the bottom of the centrifuge conical tube

**Note:** At this step (if cells are not needed, e.g., cells from culture flask #2 can be frozen) according to protocol #2

7. Add 2 ml of DMEM with 10% fetal bovine serum and dissolve the cell pellet by gently pipetting up and down.
8. Count the cells in the cell suspension using a Bürker chamber or similar. Dilute cell suspension to have 150 000 cells/mL (in DMEM + 10% FBS).

**Note:** In case of no cell counter chamber, take 400 ul of cell suspension and place it into a Petri dish or 24 well-plate, and fill up 2 ml with culture medium.

9. Prepare two culture flasks and 4 Petri dishes.
10. Take 2 mL of cell suspension into one Petri dish with an internal diameter of 34 mm (TPP Petri dish surface is 9.2 cm^2^). The cell volume (or number) has to be adjusted for a different size Petri dish. Let the cells grow in a CO2 incubator for 48h prior to AFM measurements.
11. The remaining cell suspension should be divided and put into new culture flasks (not provided). Then add culture medium (4 mL) and place culture flasks into the CO2 incubator.

**Note:** The passage ratio for these cells is: 1:2 or 1:4 (we recommend using 1:3).

**Protocol #2 (freezing cells):**

7. Conduct steps 1 to 6 from procedure #1.
8. Add 3 mL of freezing medium (10% DMSO dissolved in culture medium, here DMEM), mix, and aliquot the cell suspension into a cryogenic vial (typically, 1 mL per one vial).
9. Vials must be frozen at -80°C for 4 hours and, later on, placed in liquid nitrogen.

**Note:** Own protocol to freeze cells can be applied, too.

#### 3. Cell Preparation for AFM measurements

1. Take one Petri dish prepared for AFM measurements (see section 2, protocol #1)
2. Exchange medium with a fresh culture medium containing 10 mM of HEPES.

**Note:** Take 10 mL of culture medium (DMEM + 10% FBS +1 % antibiotics) and add 100 uL of 1 M HEPES to reach the concentration of 10 mM) – use this solution in the AFM measurements.

#### 4. Calibration of the AFM

**Note:** The proposed AFM measurements were conducted at 37°C. For room temperature measurements, adapt accordingly.

**Note:** The cantilevers chosen for the study were hemispherical MLCT-SPH-DC E, precalibrated by the manufacturer. Otherwise, it is advisable to use spherical probes with a spring constant above 0.1 N/m.

2. Document the experiment:

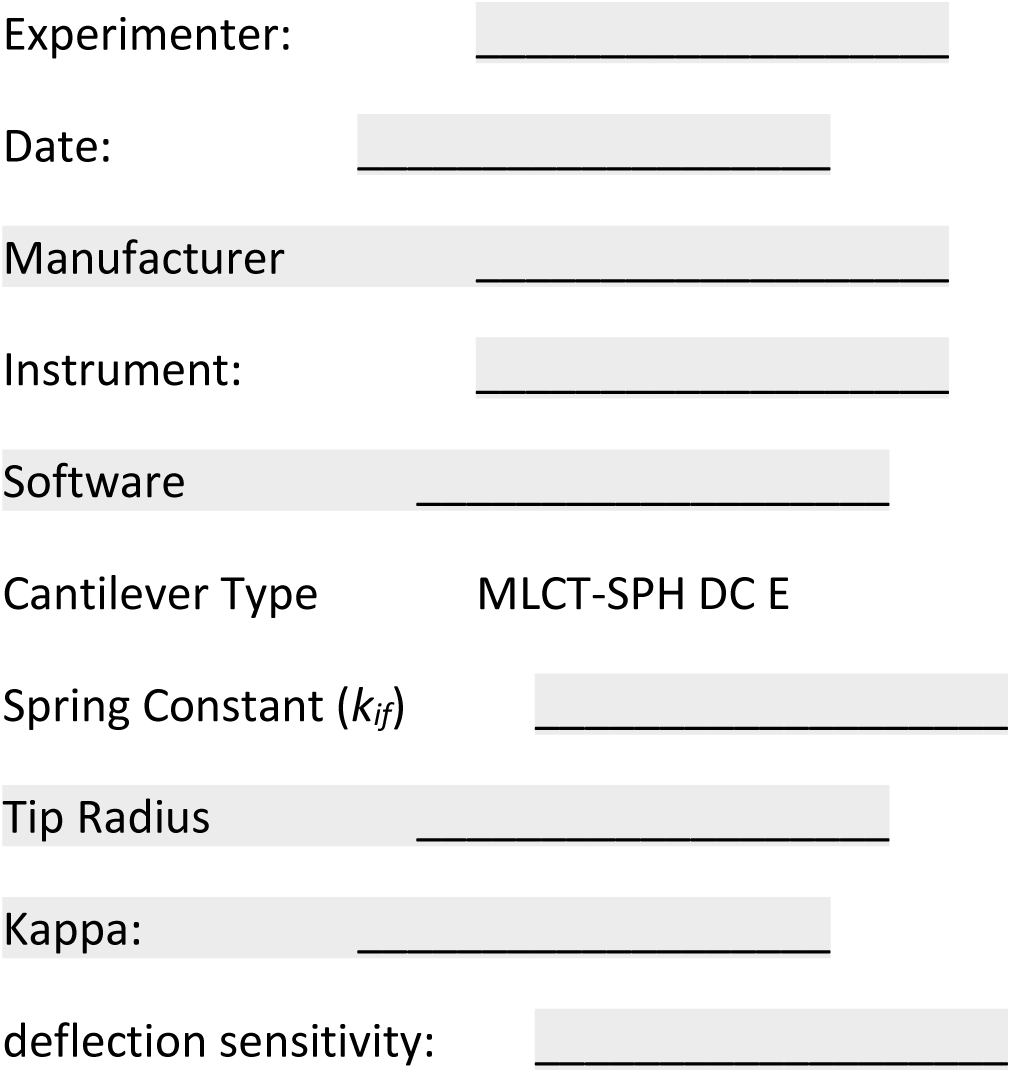

3. Apply the SNAP procedure for deflection sensitivity calibration

**Note:** SNAP procedure is described in an open-access journal *Scientific Reports* (Schillers et al. *Scientific Reports* 7 (2017) 5117.

4. Set the kappa factor to a value of 1.11.

**Note:** The kappa factor is the ratio between the freely oscillating cantilever (as in a thermal) and the cantilever in contact (as in a force curve).

5. Mount the cantilever and the sample in the AFM. Immerse the cantilever in the culture medium, adjust the laser beam to the cantilever probe, and adjust the photo-detector position.

**Note:** A bare Petri dish (without cells) to calibrate without time pressure is recommended.

6. Calibration deflection sensitivity *S* is the most critical part of setting up the instrument. Thermals should be of high quality; thus, sample and average data for at least 20s are recommended. There are two ways of deflection sensitivity calibration, dependent on the AFM instrument type:

i. If the AFM instrument allows for a non-touch calibration, which requires taking a thermal, enter the force spring constant (precalibrated value given by the manufacturer) and let the software calculate the deflection sensitivity.
ii. If the AFM instrument does not allow this procedure, the following procedure can be applied:
- type in a reasonable deflection sensitivity
- record a thermal, analyze it to get a force constant *kth*
- change the deflection sensitivity by multiplying it with: 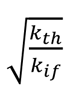
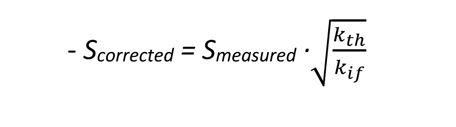

**Note:** After calibration is done, save the last recorded thermal, export it as text, and take a screenshot of the thermal in case of trouble with its importing.

7. Document the obtained deflection sensitivity: Filename of thermal:

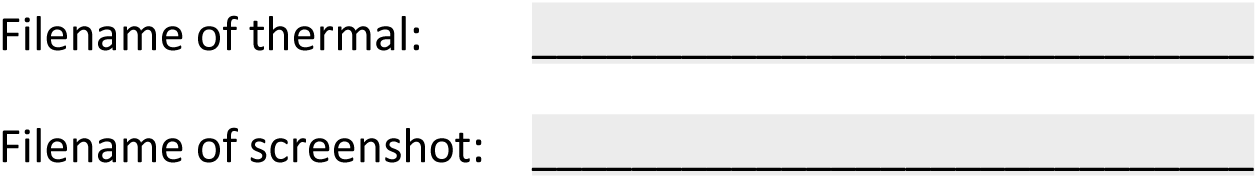

### **5.** AFM measurements on cells

1. Move the cantilever above a center of a cell.
2. Take an optical image of the place chosen to be measured.
3. Measure 10 different locations separated by at least 200 microns. Locate cellular islets/regions with the features indicated in the attached optical image.

**Note:** If the maximum time of 2 hours is exceeded, change the sample (use another Petri dish)

**Note:** Choose flat cells as it is indicated in the image of PANC-1 cells:

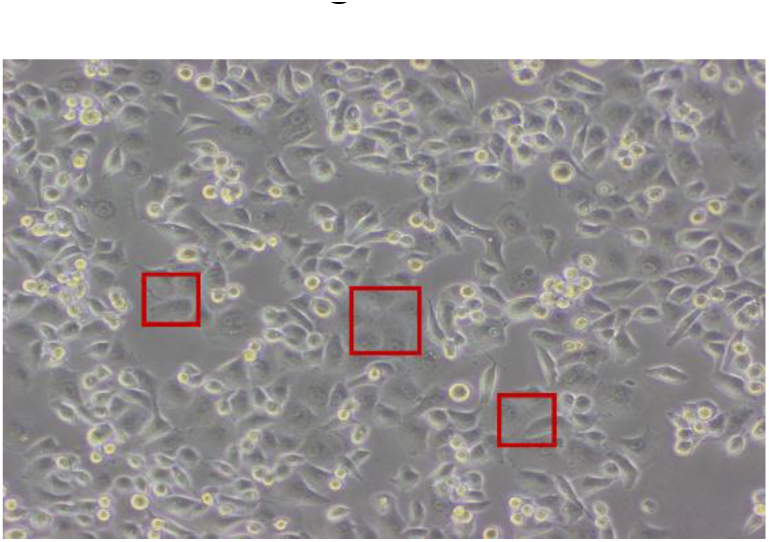

4. Record force volumes with the following parameters:

z-travel distance: 5 µm

number of points in the force curve: possibly 1 point/nm (i.e. n > 5000) tip velocity: 5µm/s

force scan rate 0.5 curves per second

trigger point (the point at which approach ends): 700 nm or 7 nN number of curves per volume 10 curves x 10 curves

area of force volume 50 µm

z-close loop: ON, or alternatively also save the z-sensor channel

5. Document the force volume collected:

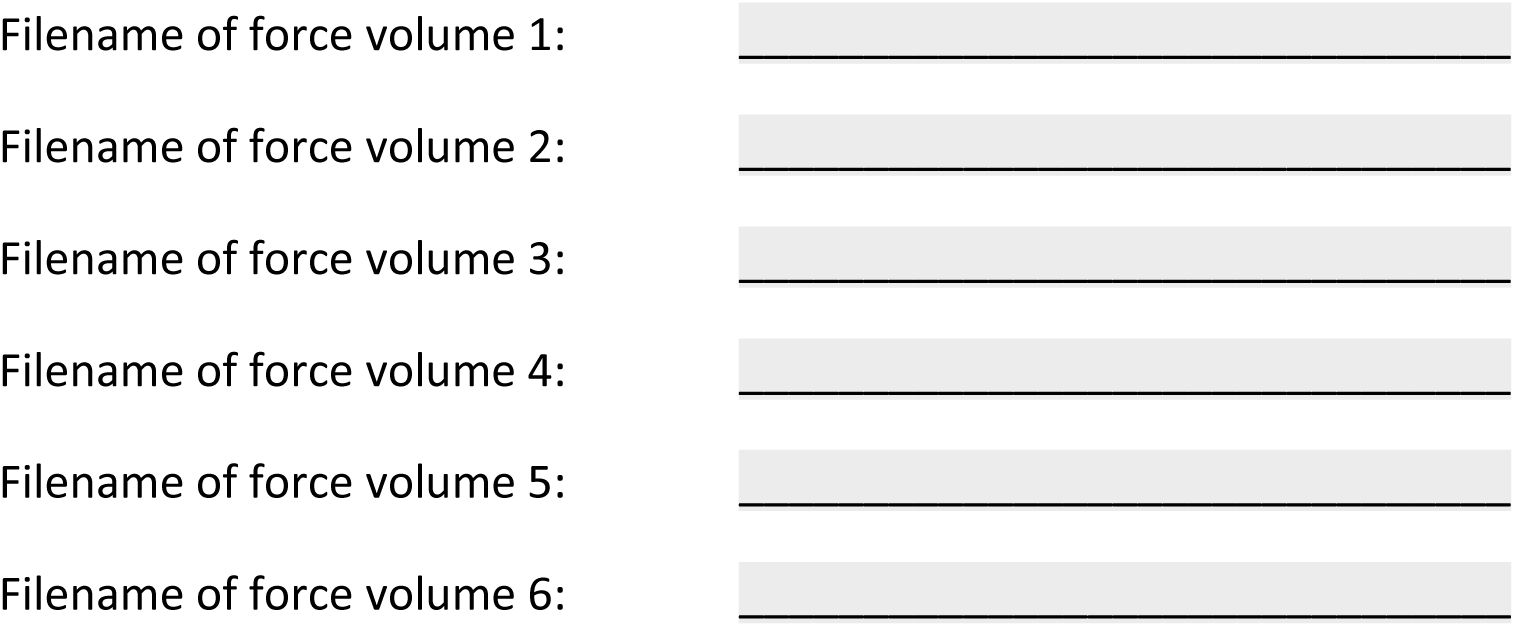
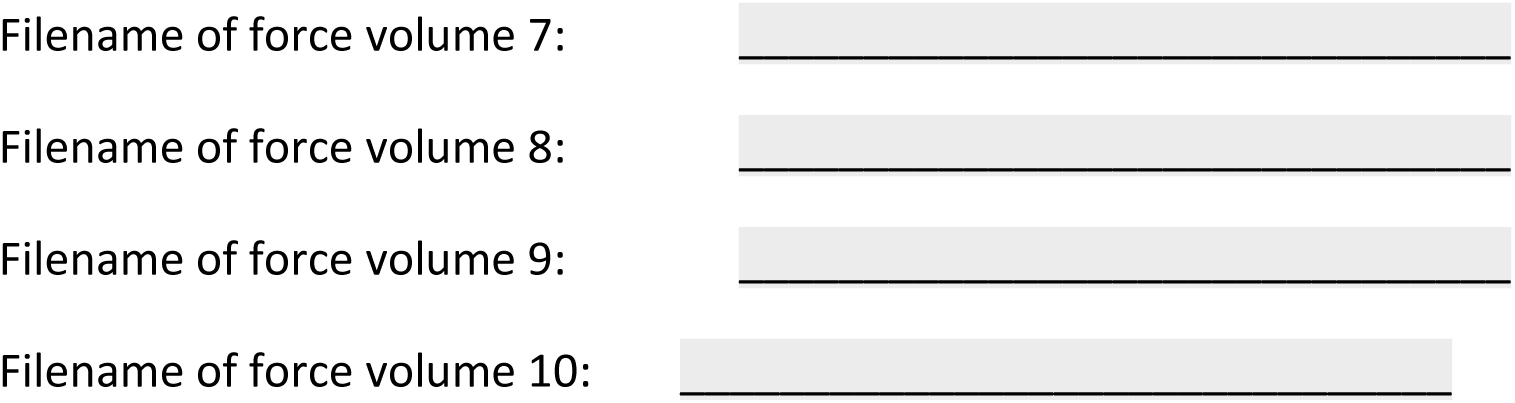

**6.** Analysing force data using local software

1. Analyze force volume data using the Hertz model for a spherical (parabolic) tip by fitting the whole data range.
2. Use the radius of the probing sphere provided by the manufacturer with the cantilevers.
3. Use a Poisson ratio of 0.5 and disable tilt correction.
4. Calculate for each force volume the median and standard deviation of elastic modulus values: Force volume 1:avg: Pa sdv: Pa

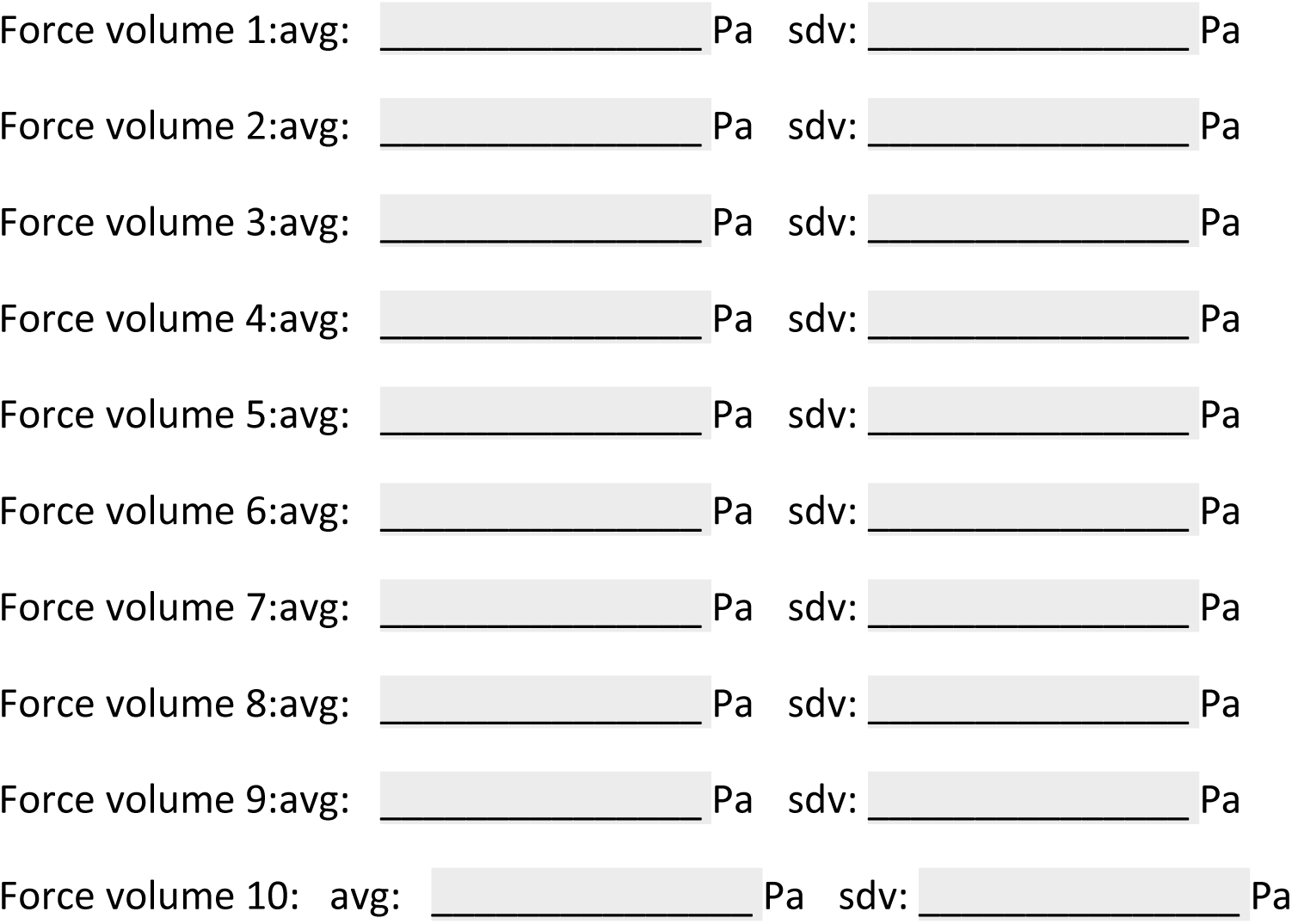

## Notes and references

1 G. Binnig, C. F. Quate and C. Gerber, Phys. Rev. Lett.,1986, 56, 930–933.

2 A. L. Weisenhorn, M. Khorsandi, S. Kasas, V. Gotzos and H. J. Butt, Nanotechnology, 1993, 4, 106–113.

3 M. Radmacher, M. Fritz and P. K. Hansma, Biophys. J., 1995, 69, 264–270.

4 W. H. Goldmann and R. M. Ezzell, Exp. Cell Res., 1996, 226, 234–237.

5 M. Lekka, P. Laidler, D. Gil, J. Lekki, Z. Stachura and A. Z. Hrynkiewicz, Eur. Biophys. J., 1999, 28, 312–316.

6 Q. S. Li, G. Y. H. Lee, C. N. Ong and C. T. Lim, Biochem. Biophys. Res. Commun., 2008, 374, 609–613.

7 E. C. Faria, N. Ma, E. Gazi, P. Gardner, M. Brown, N. W. Clarke and R. D. Snook, Analyst, 2008, 133, 1498–1500.

8 W. Xu, R. Mezencev, B. Kim, L. Wang, J. McDonald and T. Sulchek, PLoS One, 2012, e46609.

9 M. Prabhune, G. Belge, A. Dotzauer, J. Bullerdiek and M. Radmacher, Micron, 43, 1267–1272.

10 A. V. A. H. Nguyen, K. D. Nyberg, M. B. Scott, A. M. Welsh, A. V. A. H. Nguyen, N. Wu, S. V. Hohlbauch, N. A. Geisse, E. A. Gibb, A. G. Robertson, T. R. Donahue and A. C. Rowat, Integr. Biol. (United Kingdom), 2016, 8, 1232–1245.

11 X. Guo, K. Bonin, K. Scarpinato and M. Guthold, New J. Phys., DOI:10.1088/1367-2630/16/10/105002.

12 X. Tang, T. B. Kuhlenschmidt, Q. Li, S. Ali, S. Lezmi, H. Chen, M. Pires-Alves, W. W. Laegreid, T. A. Saif and M. S. Kuhlenschmidt, Mol. Cancer, , DOI:10.1186/1476-4598-13-131.

13 K. Pogoda, L. Chin, P. C. Georges, F. J. Byfield, R. Bucki, R. Kim, M. Weaver, R. G. Wells, C. Marcinkiewicz and P. A. Janmey, New J. Phys., 2014, 16, 075002.

14 M. Y. M. Chiang, Y. Yangben, N. J. Lin, J. L. Zhong and L. Yang, Biomaterials, 2013, 34, 9754– 9762.

15 H. Hertz, J. fur die Reine und Angew. Math., 1882, 1882, 156–171.

16 I. N. Sneddon, Int. J. Eng. Sci., 1965, 3, 47–57.

17 A. B. Mathur, A. M. Collinsworth, W. M. Reichert, W. E. Kraus and G. A. Truskey, J. Biomech., 2001, 34, 1545–1553.

18 J. Zemła, J. Bobrowska, A. Kubiak, T. Zieliński, J. Pabijan, K. Pogoda, P. Bobrowski and M. Lekka, Eur. Biophys. J., 2020, 49, 485–495.

19 A. Cartagena and A. Raman, Biophys. J., 2014, 106, 1033–1043.

20 X. Wang, R. Bleher, L. Wang, J. G. N. Garcia, S. M. Dudek, G. S. Shekhawat and V. P. Dravid, Sci. Rep., , DOI:10.1038/s41598-017-14722-0.

21 H. Schillers, C. Rianna, J. Schäpe, T. Luque, H. Doschke, M. Wälte, J. J. Uriarte, N. Campillo, G. P. A. Michanetzis, J. Bobrowska, A. Dumitru, E. T. Herruzo, S. Bovio, P. Parot, M. Galluzzi, A. Podestà, L. Puricelli, S. Scheuring, Y. Missirlis, R. Garcia, M. Odorico, J. M. Teulon, F. Lafont, M. Lekka, F. Rico, A. Rigato, J. L. Pellequer, H. Oberleithner, D. Navajas and M. Radmacher, Sci. Rep., 2017, 7, 5117.

22 M. Lekka, D. Gil, K. Pogoda, J. Dulińska-Litewka, R. Jach, J. Gostek, O. Klymenko, S. Prauzner-Bechcicki, Z. Stachura, J. Wiltowska-Zuber, K. Okoń and P. Laidler, Arch. Biochem. Biophys., 2012, 518, 151–156.

23 L. M. Rebelo, J. S. De Sousa, J. Mendes Filho and M. Radmacher, Nanotechnology, 2013, 24, 055102.

24 J. R. Ramos, J. Pabijan, R. Garcia and M. Lekka, BEILSTEIN J. Nanotechnol., 2014, 5, 447–457.

25 A. Calzado-Martín, M. Encinar, J. Tamayo, M. Calleja and A. San Paulo, ACS Nano, 2016, 10, 3365–3374.

26 A. Kubiak, M. Chighizola, C. Schulte, N. Bryniarska, J. Wesolowska, M. Pudelek, M. Lasota, D. Ryszawy, A. Basta-Kaim, P. Laidler, A. Podestà and M. Lekka, Nanoscale, 2021, 13, 6212–6226.

27 E. K. Dimitriadis, F. Horkay, J. Maresca, B. Kachar and R. S. Chadwick, Biophys. J., 2002, 82, 2798–2810.

28 L. Puricelli, M. Galluzzi, C. Schulte, A. Podestà and P. Milani, Rev. Sci. Instrum., 2015, 86, 033705.

29 P. D. Garcia and R. Garcia, Biophys. J., 2018, 114, 2923–2932.

30 P. D. Garcia and R. Garcia, Nanoscale, 2018, 10, 19799–19809.

31 N. Gavara and R. S. Chadwick, Nat. Nanotechnol., 2012, 7, 733–736.

32 S. Iyer, R. M. Gaikwad, V. Subba-Rao, C. D. Woodworth and I. Sokolov, Nat. Nanotechnol., 2009, 4, 389–393.

33 S. V. Beekmans and D. Iannuzzi, Surf. Topogr. Metrol. Prop., , DOI:10.1088/2051-672X/3/2/025004.

34 M. M. Lekka, K. Pogoda, J. Gostek, O. Klymenko, S. Prauzner-Bechcicki, J. Wiltowska-Zuber, J. Jaczewska, J. Lekki and Z. Stachura, MICRON, 2012, 43, 1259–1266.

35. S. Pérez-Domínguez, S. G. Kulkarni, C. Rianna and M. Radmacher, 2020.

36 C. Rianna and M. Radmacher, Nanoscale, 2017, 9, 11222–11230.

