## Supporting Materials with Annex 1 for "Reliable, standardized measurements for cell mechanical properties"

**List S1.**

lab 1 – Nanowizard 3.0 (Bruker-JPK)

lab 2 – Nanowizard 4.0 (Bruker-JPK)

lab 3 – Nanowizard 4.0 (Bruker-JPK)

lab 4 – Pavone, Piuma (Optics 11)

lab 5 – MFP3D (Asylum Research, Santa Barbara, CA, USA)

lab 6 – Bioscope Catalyst (Bruker Nano, Santa Barbara/CA, USA)

lab 7 – BioScope Resolve (Bruker Nano, Santa Barbara/CA, USA)

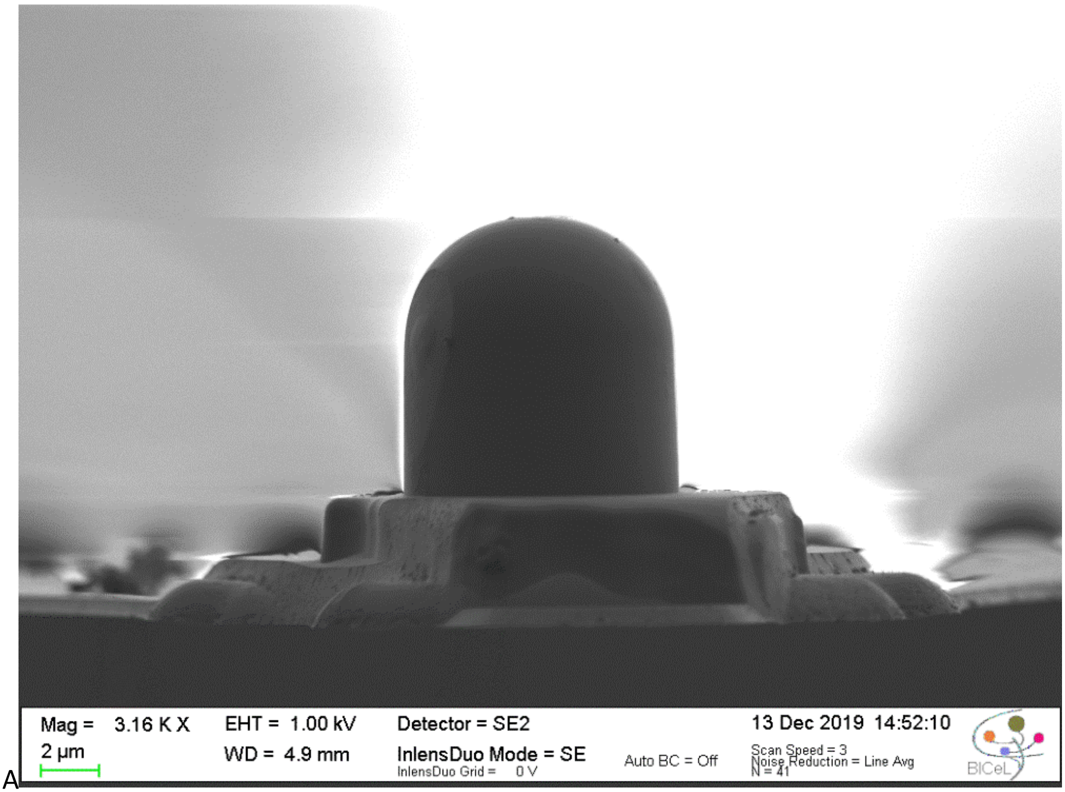

A

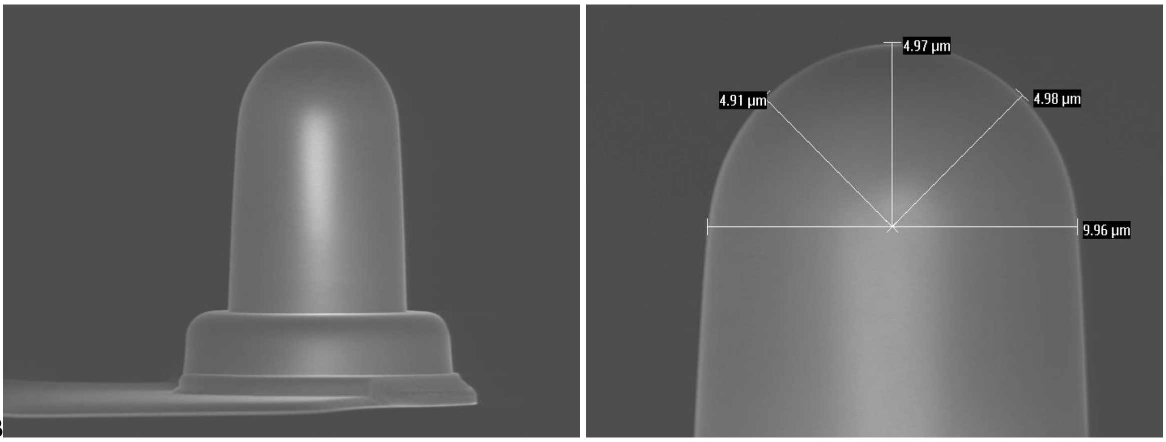

B

**Supporting Figure S1. A)** SEM (Scanning Electron Microscope) image of the MLCT-SPH-DC silicon nitride cantilevers (Bruker) with a hemispherical tip. The image was collected using SEM available in BICeL facility (Lille, France). B) SEM images of the currently available cantilevers with SPH tips available for AFM community (courtesy of A.Dulebo, Bruker).

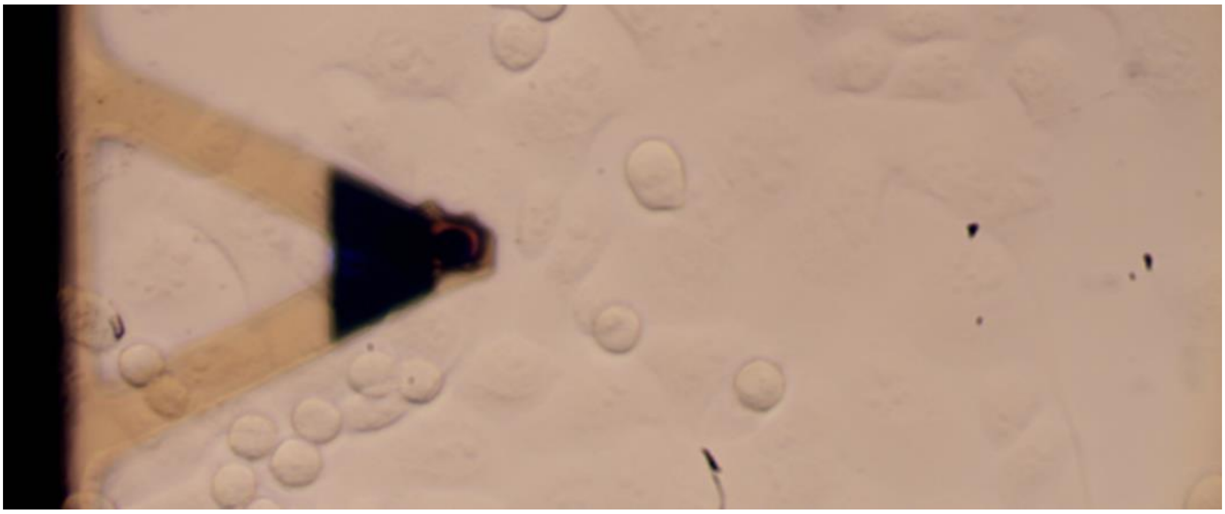

**Supporting Figure S2.** Top view image of the PANC-1 monolayer, the flat area of the monolayer measured in standardization experiments. Image was collected with the top view optics integrated with MFP3D (Asylum Research, Santa Barbara, CA, USA).

### Annex 1: Protocol for measuring cell mechanics

*Sandra Pérez-Domínguez[1], Shruti G. Kulkarni[1], Joanna Pabijan[1], Kajangi Gnanachandran[2], Hatice Holuigue[3], Mar Eroles[6], Ewelina Lorenc[3], Massimiliano Berardi[4,5], Nelda Antonovaite[5], Maria Luisa Marini[7], Javier Lopez Alonso[7], Lorena Redonto-Morata[7], Vincent Dupres[7], Sebastien Janel[7], Sovon Acharya[8], Jorge Otero[9], Daniel Navajas[9], Kevin Bielawski[5], Hermann Schillers[8], Frank Lafont[7], Felix Rico[6], Alessandro Podestà[*3], Manfred Radmacher[*1], Małgorzata Lekka[*2]*

[1] Institute of Biophysics, University of Bremen, 28359, Bremen, Germany

[2] Department of Biophysical Microstructures, Institute of Nuclear Physics, Polish Academy of Sciences, PL-31342 Kraków, Poland

[3] Department of Physics "Aldo Pontremoli" and CIMAINA, University of Milano, via Celoria 16, 20133 Milano, Italy,

[4] Laserlab, Department of Physics and Astronomy, Vrije Universiteit Amsterdam, De Boelelaan 1081, 1081 HV, Amsterdam, The Netherlands

[5] Optics11 life, Hettenheuvelweg 37-39, 1101 BM, Amsterdam, The Netherlands

[6] Aix-Marseille Univ, CNRS, INSERM, LAI, Turing centre for living systems, Marseille, France

[7] Université de Lille, CNRS, INSERM, CHU Lille, Institut Pasteur de Lille, U1019-UMR9017, CIIL—Center for Infection and Immunity of Lille, F-59000 Lille, France

[8] Institute of Physiology II, University Muenster, Robert-Koch-Str. 27b, 48149 Muenster, Germany

[9] Institute for Bioengineering of Catalonia and Universitat de Barcelona, Barcelona, Spain. CIBER de Enfermedades Respiratorias, Madrid, Spain.

**Protocol sheet for measuring cell mechanics**

*TOC*

0. General Information

1. Shipment of cells

2. Cell cultures

3. Preparing cells for AFM measurements

4. Calibration of the AFM

5. AFM measurements on cells

6. Analysing force data.

7. Additional info on cells

**0. General Information**

Materials needed:

- two 25 $cm^2$ culture flasks (#1, #2) with living cells (here, the protocol was tested for pancreatic cancer PANC-1 cells);

**Note:** PANC-1 cells were established from a human pancreatic cancer, isolated from a pancreatic carcinoma of ductal cell origin of a 56-year-old male (Lieber et al. International Journal of Cancer. 15 (1975) 741–747, doi:10.1002/ijc.2910150505). These cells have an epithelial morphology and are adherent in cell culture flasks; however, they tend to clump. PANC-1 cells can metastasize, but they have poor differentiation abilities.

- four empty culture flasks (sterile)
- two 50 ml tubes of culture medium (DMEM + 10% FBS) with antibiotics (1%) - (sterile);
- one 50 ml tube of PBS (sterile)
- one 15 ml tube with trypsin solution (sterile);
- one 15 ml tube of freezing medium: DMEM + 5 % DMSO (sterile)
- one 2 ml tube of 1 M HEPES (sterile)

Cell Culture Dish, 35x10mm – TPP cat no. *93040*
Tissue Culture Flasks 25 cm2 – TPP cat no. 90025

DMEM - ATCC cat no. 30-2002
FBS - SIGMA cat no. F9665
Trypsin-EDTA solution (10x) - SIGMA cat no. T4174
Dimethyl sulfoxide (DMSO) – SIGMA cat no. D2438
HEPES buffer (1 M) – SIGMA cat no. H0887
PBS – SIGMA cat no. P4417
Antibiotics (Penicillin − Streptomycin − Neomycin Solution Stabilized) – SIGMA, cat no P4083

**1. Shipment of cells**

Living cells can be shipped at 50-60% confluency; culture flasks should be filled with a maximum amount of culture medium (for PANC-1 cells: DMEM + 10% FBS + 1% antibiotics).

1. Document sample and shipping details:

Culture flask number: ______

Prepared by: ____________________

Preparation date: ____________________

Sent data: ____________________

Sent to (location) ____________________

Arrival date & time: ____________________

Flask 1 Passaging date & time ____________________

Flask 2 Passaging date & time ____________________

**2. Cell culture**

Work under sterile conditions (under the laminar flow chamber):

Experimenter: ____________________

Date: ____________________

Manufacturer ____________________

Instrument: ____________________

Software ____________________

Cantilever Type MLCT-SPH DC E

Spring Constant ($k_{if}$) ____________________

Tip Radius ____________________

Kappa: ____________________

deflection sensitivity: ____________________

3. Apply the SNAP procedure for deflection sensitivity calibration

**Note:** SNAP procedure is described in an open-access journal *Scientific Reports* (Schillers et al. *Scientific Reports* 7 (2017) 5117.

4. Set the kappa factor to a value of 1.11.

**Note:** The kappa factor is the ratio between the freely oscillating cantilever (as in a thermal) and the cantilever in contact (as in a force curve).

5. Mount the cantilever and the sample in the AFM. Immerse the cantilever in the culture medium, adjust the laser beam to the cantilever probe, and adjust the photo-detector position.

**Note:** A bare Petri dish (without cells) to calibrate without time pressure is recommended.

6. Calibration deflection sensitivity *S* is the most critical part of setting up the instrument. Thermals should be of high quality; thus, sample and average data for at least 20s are recommended. There are two ways of deflection sensitivity calibration, dependent on the AFM instrument type:
   i. If the AFM instrument allows for a non-touch calibration, which requires taking a thermal, enter the force spring constant (precalibrated value given by the manufacturer) and let the software calculate the deflection sensitivity.
   ii. If the AFM instrument does not allow this procedure, the following procedure can be applied:

   - type in a reasonable deflection sensitivity

   - record a thermal, analyze it to get a force constant $k_{th}$

   - change the deflection sensitivity by multiplying it with: $\sqrt{\frac{k_{th}}{k_{if}}}$

   - $S_{corrected} = S_{measured} \cdot \sqrt{\frac{k_{th}}{k_{if}}}$

**Note:** After calibration is done, save the last recorded thermal, export it as text, and take a screenshot of the thermal in case of trouble with its importing.

7. Document the obtained deflection sensitivity:

   Filename of thermal: ________________________________

   Filename of screenshot: ________________________________

**5. AFM measurements on cells**

1. Move the cantilever above a center of a cell.
2. Take an optical image of the place chosen to be measured.
3. Measure 10 different locations separated by at least 200 microns. Locate cellular islets/regions with the features indicated in the attached optical image.

**Note:** If the maximum time of 2 hours is exceeded, change the sample (use another Petri dish)

**Note:** Choose flat cells as it is indicated in the image of PANC-1 cells:

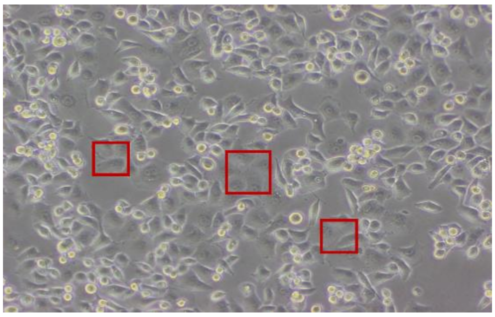

4. Record force volumes with the following parameters:
   z-travel distance: 5 µm

   number of points in the force curve: possibly 1 point/nm (i.e. n > 5000)

   tip velocity: 5µm/s

   force scan rate 0.5 curves per second

   trigger point (the point at which approach ends): 700 nm or 7 nN

   number of curves per volume 10 curves x 10 curves

   area of force volume 50 µm

   z-close loop: ON, or alternatively also save the z-sensor channel

5. Document the force volume collected:
   Filename of force volume 1: ___________________________________

   Filename of force volume 2: ___________________________________

   Filename of force volume 3: ___________________________________

   Filename of force volume 4: ___________________________________

   Filename of force volume 5: ___________________________________

   Filename of force volume 6: ___________________________________

Filename of force volume 7: ________________________________

Filename of force volume 8: ________________________________

Filename of force volume 9: ________________________________

Filename of force volume 10: ________________________________

**6. Analysing force data using local software**

1. Analyze force volume data using the Hertz model for a spherical (parabolic) tip by fitting the whole data range.
2. Use the radius of the probing sphere provided by the manufacturer with the cantilevers.
3. Use a Poisson ratio of 0.5 and disable tilt correction.
4. Calculate for each force volume the median and standard deviation of elastic modulus values:

Force volume 1:avg: ________________ Pa sdv: ________________ Pa

Force volume 2:avg: ________________ Pa sdv: ________________ Pa

Force volume 3:avg: ________________ Pa sdv: ________________ Pa

Force volume 4:avg: ________________ Pa sdv: ________________ Pa

Force volume 5:avg: ________________ Pa sdv: ________________ Pa

Force volume 6:avg: ________________ Pa sdv: ________________ Pa

Force volume 7:avg: ________________ Pa sdv: ________________ Pa

Force volume 8:avg: ________________ Pa sdv: ________________ Pa

Force volume 9:avg: ________________ Pa sdv: ________________ Pa

Force volume 10: avg: ________________ Pa sdv: ________________ Pa
